# Resource-efficiency of cyanobacterium production on Mars: Assessment and paths forward

**DOI:** 10.1101/2024.07.29.605555

**Authors:** Tiago P. Ramalho, Vincent Baumgartner, Nils Kunst, David Rodrigues, Emma Bohuon, Basile Leroy, Guillaume Pillot, Christiane Heinicke, Sven Kerzenmacher, Marc Avila, Cyprien Verseux

## Abstract

Space agencies and private companies strive for a permanent human presence on the Moon and ultimately on Mars. Bioprocesses have been advocated as key enablers due to their ability to transform locally available resources into added-value materials. However, the resource-efficiency and scaling of space biosystems remain poorly understood, hindering quantitative estimates of their potential performance. We leveraged extensive cultivation experiments, where a cyanobacterium (*Anabaena* sp. PCC 7938) was subject to conditions attainable on Mars, to develop a model that can estimate bioprocess productivity and resource-efficiency as a function of water, light, temperature, regolith minerals and perchlorates, and atmospheric carbon and nitrogen. We show that a breakeven can be reached within a few years. We discuss research lines to improve both resource efficiency and the accuracy of the model, thereby reducing the need for costly tests in space and eventually leading to a biotechnology-supported, sustained human presence on Mars.

**TEASER:** Bioprocess modeling shows that cyanobacterium-based biotechnologies can be a sustainable basis for resource production on Mars.

## INTRODUCTION

The world’s leading space agencies share the goal of sending humans to Mars within the coming two decades (*1*–*3*). Such crewed missions could lead to a wealth of scientific discoveries beyond the reach of robots alone, pertaining for instance to geosciences, human physiology, fundamental physics or the search for life beyond Earth (*4, 5*). Holding this promise requires a sustainable program: Mars exploration should not end with early missions, no more than the exploration of Antarctica stopped with its Heroic Age. A permanent presence is warranted, perhaps analogous to that in today’s polar stations. However, the costs needed to haul payloads from Earth make supplying the consumables required for human survival a challenge. Most of these consumables should be produced on site (*6*).

Bioprocesses are particularly suited to this task: a wide range of resources as critical as food, fuel, pharmaceuticals or structural materials could be produced and recycled through them (*7*–*10*). One difficulty is that the bulk of the required feedstock should not be imported from Earth: while recycling could minimize the need for resupplies (*11*), it would not eliminate it entirely since recycling efficiencies are limited. A promising solution lies in the utilization of materials available on site, a strategy referred to as “in situ resource utilization” (ISRU). But most organisms of biotechnological interest likely cannot use raw Martian resources (*12*), which are – to current knowledge – poor in organics, fixed nitrogen and readily available mineral nutrients.

One strategy proposed to feed these organisms lies in using selected species of diazotrophic, rock-weathering cyanobacteria as primary producers (*12*). An expanding body of work suggests that these could be fed with local resources: water mined from the ground and atmosphere; carbon and nitrogen sourced from the atmosphere as CO_2_ and N_2_ (*13, 14*); and other nutrients leached from the local regolith (the Martian soil) (*15*–*17*). The cultivated cyanobacteria could then produce various consumables directly, such as O_2,_ dietary proteins or biomaterials (*18, 19*), but also be used for feeding heterotrophic microorganisms (*13, 20, 21*), plants (*22*), or other secondary producers. So far, however, the resource-efficiency of such a process – as well as how that resource-efficiency could be improved – has remained poorly understood. It is this deficiency which we address here.

We describe a mathematical model which, fed with input values from literature and some biological parameters determined in the present study, enables assessments of productivity and resource-efficiency of cyanobacterium cultivation on Mars through ISRU. We also present results obtained using this model, which suggest the viability of such cultivation: despite conservative assumptions, we predict a breakeven within five years of operation. Although these results are not definitive – we were limited by the many uncertainties that remain, notably on technologies that will become available and on specific mission parameters –, they do provide insights which can guide future development efforts.

## RESULTS

Based on the mathematical model we developed (see “Methods”), we assessed the productivity and resource-efficiency of a cyanobacterium cultivation system on Mars that relies on ISRU. Unless specified otherwise, the results presented below assume a set of parameters (collectively referred to as the “Main scenario”; see Table 1) deemed most realistic given today’s level of advancement. In this scenario, the cyanobacterium *Anabaena* sp. PCC 7938 is grown planktonically in an airlift photobioreactor maintained at an optimal growth temperature (304.89 K). It is cultivated in 30-day runs, each starting with an inoculum representing a small fraction (below one percent) of the biomass eventually produced. Mineral nutrients come exclusively from windblown regolith containing 0.4 wt% of perchlorates and provided at a concentration of 200 kg m^-3^. The culture is continuously illuminated with up to 650 mol_ph_ m^-2^ s^-1^ at the reactor surface. The impact of deviations from these parameter values are also presented, individually or combined into alternative scenarios.

**Table 1.** Main and alternative scenarios. Symbols in the “Inputs” column are explained in Table S1.

| Scenario | Inputs | Description |
| --- | --- | --- |
| Main | Main operational inputs: $T = 304.89 \text{ K}$; $H_{\text{culture}} = 1.7 \text{ m}$; $v_{\text{gsmax}} = 0.08 \text{ m s}^{-1}$; $\text{wt}\% = 0.4$; $\text{BET} = 7605.4 \text{ m}^2 \text{ kg}^{-1}$; $C_R = 200 \text{ kg m}^{-3}$; $I_{\text{ph0}} = 6.5 \times 10^{-4} \text{ mol}_{\text{ph}} \text{ m}^{-2} \text{ s}^{-1}$; $\text{time} = 30 \text{ days}$; $C_{x0} = 0.621 \text{ mol m}^{-3}$; $C_{p0} = 0 \text{ mol m}^{-3}$. | Realistic design based on today’s state of the art, accounting for resources expected to be available on Mars and productivity requirements. |
| High maintenance | Main operational inputs; increase in yearly crew time from 5 to 20 hours; yearly additional mass cost corresponding to 50% of the initial structural mass. | Multiple reactor malfunctions and high system wear. |
| Biofilm | Main operational inputs; reduction in water usage (by 90%), structural mass (by 14%), and power and thermal control requirements of the pumps (by 92%) (20). | Cyanobacteria cultivated as biofilms instead of planktonic cells. |
| Gas ISRU | Main operational inputs except for: $C_R = 0 \text{ kg m}^{-3}$; $C_{p0} = 0.2 \text{ mol m}^{-3}$. Mass of imported salts included in the payload. | Nutrients provided from the regolith in the Main scenario are here shipped from Earth. |

### Effects of parameters related to regolith and light on productivity

The productivity of cyanobacterial cultivation from Martian resources is highly dependent on the availability of light and regolith as the sources of, respectively, energy and nutrients. The impact of varying regolith concentration, together with its perchlorate mass fraction and with light intensity, is illustrated in Figure 1.

**Figure 1.**
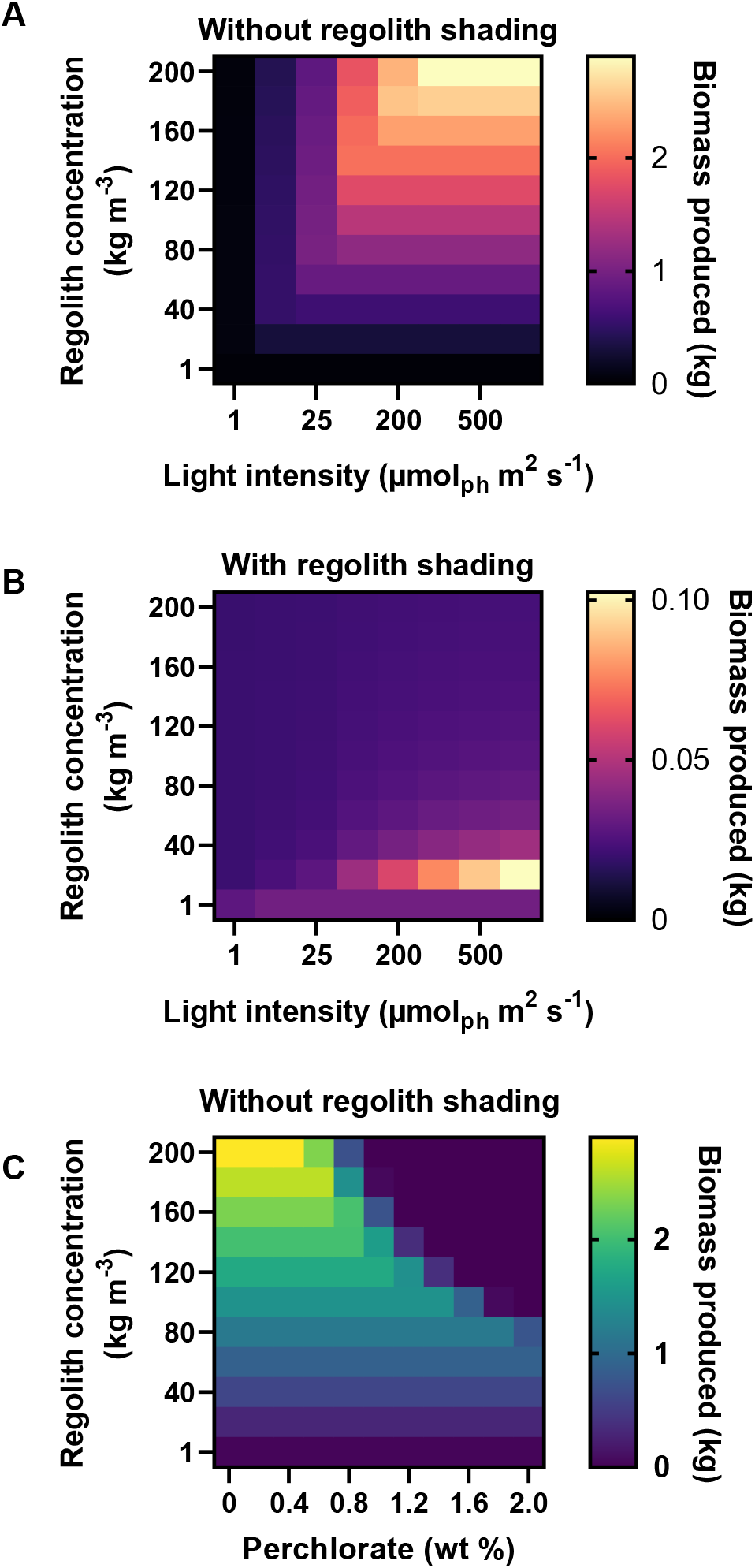
Effects of variations in parameters related to regolith and light on productivity. **A**: Biomass produced as a function of regolith concentration and light intensity. **B**: Biomass produced as a function of regolith concentration and light intensity, assuming regolith is homogeneously suspended and shades cells. **C**: Biomass produced as a function of the concentration, and perchlorate mass fraction, of the regolith. Regolith shading is assumed to be prevented.

In what otherwise corresponds to the Main scenario, productivity increases when light intensity and regolith concentration are increased jointly (either can be limiting), up to a light intensity at the reactor surface of 350 μmol_ph_ m^-2^ s^-1^ and a regolith concentration of 200 kg m^-3^ – where 2.9 kg of biomass are produced in 30 days in a one-cubic-meter reactor (Figure 1, A). This assumes that regolith grains are kept in an area of the reactor where they do not affect light penetration into the culture: if assuming that grains are homogeneously suspended in the culture medium, shading greatly reduces productivity. In this latter case, a maximum of 0.1 kg of produced biomass is reached, under a light intensity of 650 μmol_ph_ m^-2^ s^-1^ and a regolith concentration of 20 kg m^-3^.

Beside nutrients, the regolith contains oxychlorine species, including perchlorates. These seem ubiquitous at the surface (*23*) and the dependence of their concentration on depth is unknown (*24*). The direct oxidative stress they cause to bacteria in aqueous solutions is limited by a kinetic barrier (*25*). They nonetheless remain highly toxic, owing in large part to their chaotropicity (*26*). In the Main scenario, we assume a concentration of 0.4% in weight, which seems to be close to the average concentration in Rocknest windblown soil (*27*). Changes in this mass fraction affect the optimal regolith concentration, as well as the maximum production of biomass (Figure 1, C). However, this effect is notable only when the combination of regolith concentration and perchlorate mass fraction yields at least 1.2 kg m^-3^ of dissolved perchlorate. Above this value, an increase in perchlorate concentration rapidly leads to a failure to produce any biomass.

### Evolution of productivity over time

Up to a threshold, perchlorate concentrations have little impact on the biomass produced despite the effect of perchlorates on instantaneous growth rates. This is due to the dynamics of consumption (by cyanobacteria) and release (from the regolith) of the limiting nutrient. With an analogue – MGS-1 (*28*) – of the regolith type assumed in our Main scenario, this nutrient is phosphorus (*17*). To further understand the interdependency between biomass and phosphorus, their concentrations over time were estimated with the model. Following cultivation onset, phosphorus concentrations increase, reaching a maximum of 0.15 mol m^-3^ on Day 5 and supporting high growth rates (Figure 2, A). After Day 5, consumption outweighs release, causing a decline in phosphorus concentration. On Day 14, this concentration reaches its lowest value (0.003 mol m^-3^), which is maintained throughout cultivation as any released phosphorus is immediately consumed. Biomass concentration thus increases linearly until the end of the cultivation period, at a rate which is dictated by that of phosphorus release.

**Figure 2.**
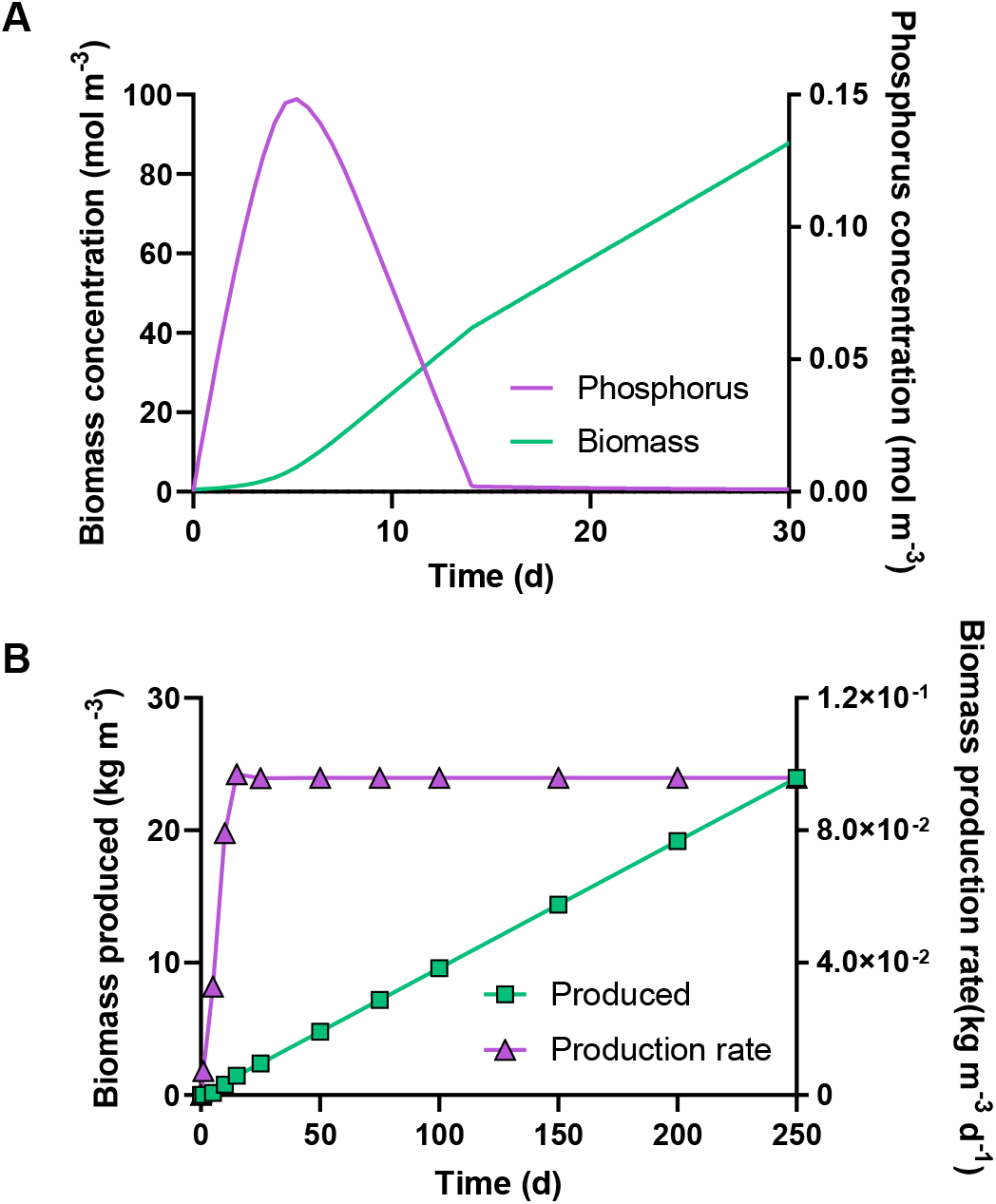
Productivity over time. **A**: Concentrations of biomass and phosphorus over 30 days of cultivation. **B**: Biomass produced, and production rate, as a function of total cultivation time and over 250 days.

More broadly, phosphorus release rates largely determine growth dynamics when regolith is limiting. The main mode of operation corresponding to the results shown in Figure 2 is a batch mode: at no point did switching to continuous cultivation increase yields. Since the duration of the batch had no impact on them either (the model does not account for a possible lag phase or the exhaustion of phosphorus in the regolith), we assumed 30-day batches in the Main scenario, as laboratory experiments have been limited to 28 days. The biomass production rates and production over a longer term (250 days) are nonetheless shown in Figure 2, B. The produced biomass increases exponentially with cultivation time until Day 5, after which it continues to accumulate linearly. These dynamics are mirrored in the biomass production rate, which quickly increases to 0.096 kg d^-1^ and hardly varies afterwards.

### Gas requirements throughout cultivation

The pressure, composition and flow rate of the gas mixture provided to the cultivation system is expected to impact resource-efficiency, notably due to the costs associated with processing the Martian atmosphere and to the mechanical constraints on hardware of a high inner pressure. The minimum partial pressures of gases required to support the growth rates predicted for the Main scenario, as well as of the gases imposed by the operating process, are shown in Figure 3, A. Their value when the total pressure is maximum is shown in Figure 3, B.

**Figure 3.**
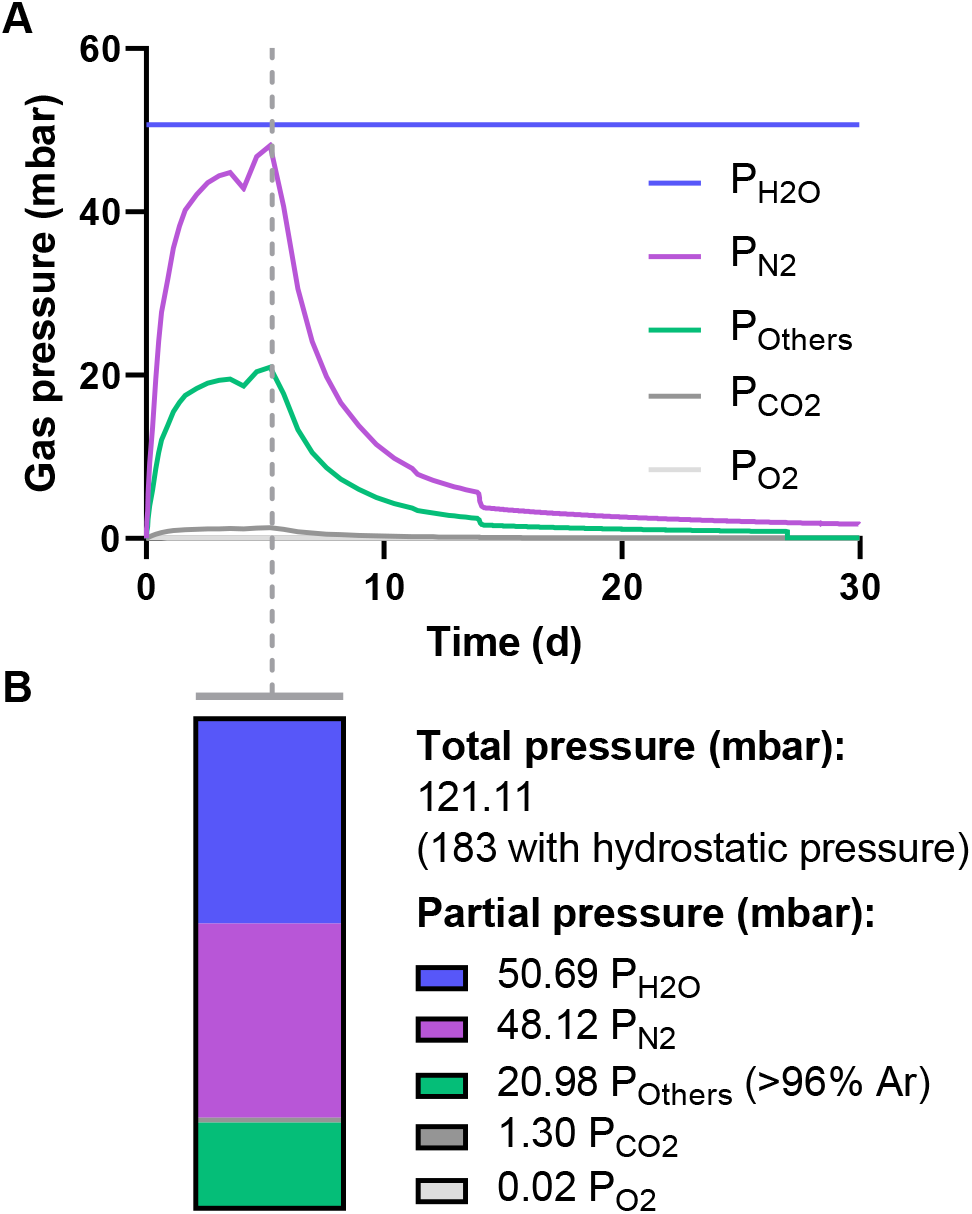
Gas requirements throughout a 30-day batch cultivation. **A**: Gas partial pressures necessary to maintain biomass productivity (these of N_2_ and CO_2_) or imposed by the operating process. **B**: Total and partial gas pressures (in mbar) when the total pressure is maximum.

The partial pressure of water vapour has a constant value of 50.69 mbar, assuming a relative humidity of 100% and including the hydrostatic pressure at the centre of the reactor. The required partial pressure of other gases varies with time. That of gaseous oxygen (a cultivation product) starts at a maximum of 0.08 mbar, then stabilizes at 0.013 mbar as the rate of its elimination through flushing matches its production rate. The partial pressures of the two gaseous nutrients – dinitrogen and carbon dioxide – peak at 48.12 and 1.30 mbar, respectively, when growth rates are maximum, but sharply decrease afterwards. Other gases from the Martian atmosphere (predominantly, argon), introduced together with dinitrogen as removing them from the dinitrogen source would cause unnecessary costs, vary similarly over time and peak at 20.98 mbar. The total pressure, on which depend the required level of robustness – and therefore the mass – of the cultivation hardware, as well as energy needs for pressurization (*13, 14*), peaks at 121.11 mbar (183 mbar with full hydrostatic pressure; Figure 3, B).

While requirements vary over time and a timely control of atmospheric conditions could improve resource-efficiency, we assume a simpler system, where atmospheric conditions are these shown in Figure 3, B, in subsequent assessments of equivalent system mass (ESM).

### Determination of equivalent system mass

The resource-efficiency of ISRU-based cyanobacterium cultivation was determined by comparing the mass of its products to its ESM – a metric for launch costs which accounts for mass, volume, power, cooling and crewtime requirements (*29*). The mass of products, the equivalent mass of spent resources (“costs”), and the ratio between both (specific ESM), over 30 years, are shown in Figure 4, A. The specific ESM at the end of the first year is 4.51 kg_eqcost_ kg_product_^-1^. Most of the costs (326.4 kg_eq_) are establishment costs: hardware mass and volume; costs related to the mining of water; and costs related to power and thermal control. Although the latter two incur over time, they are factored into establishment costs as their associated structures must be available at bioproduction onset. On a yearly basis, costs attributed to crew time and carbon dioxide acquisition (7.44 kg_eq_) are outweighed by product mass (72.33 kg), and a breakeven point (BEP) – when the specific ESM equals one – is reached within 5 years.

**Figure 4.**
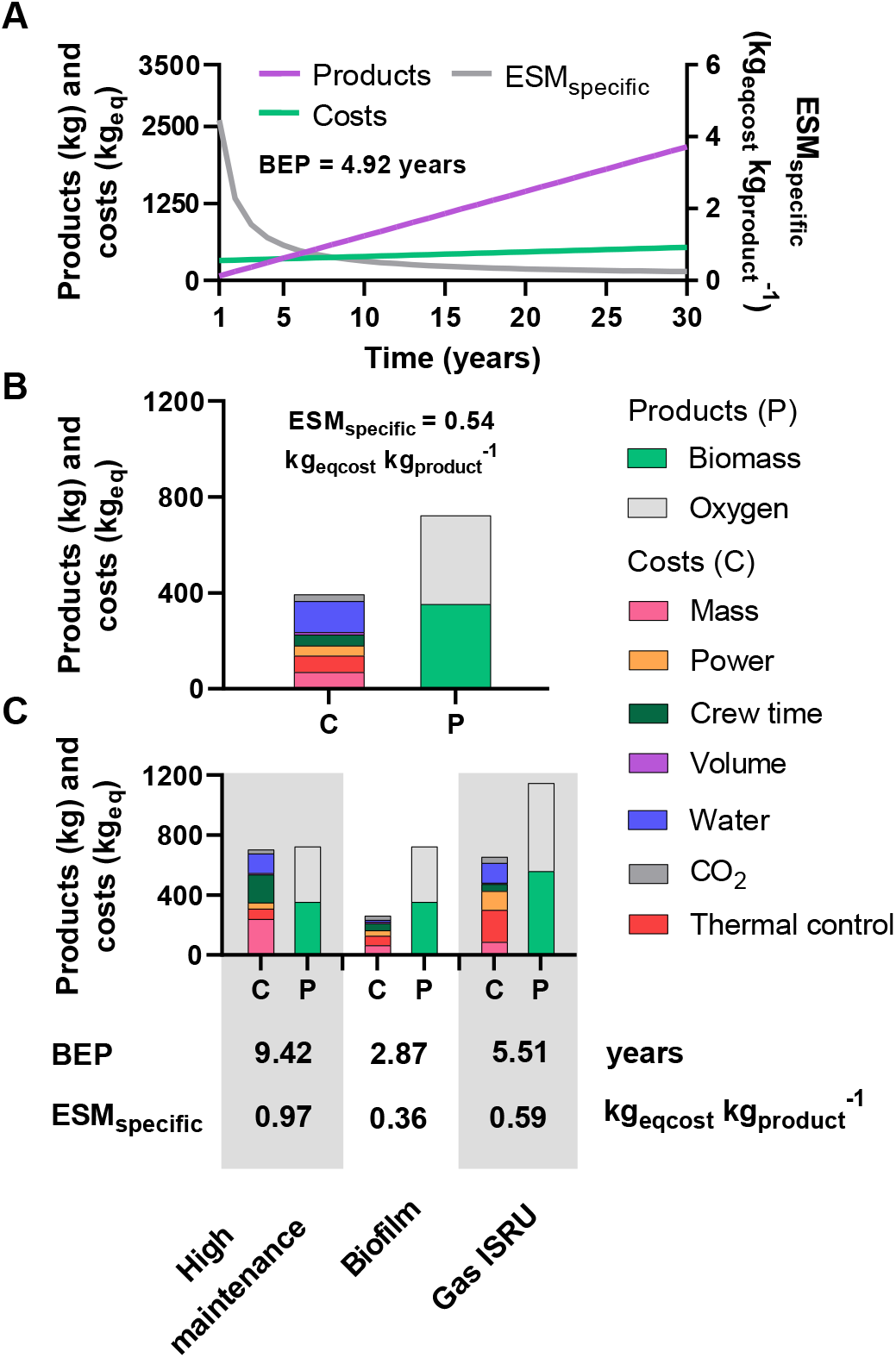
Equivalent system mass (ESM) analysis of cyanobacterium cultivation through ISRU in a one-cubic-meter photobioreactor. **A**: ESM, product mass, specific ESM (ESM_specific_) and time to reach a break-even point (BEP) over 30 years of operation in the Main scenario. **B**: Mass of products (P), costs (C), and ESM_specific_ after ten years of operation in the Main scenario. **C**: ESM analysis after ten years in alternative scenarios.

A breakdown of products and costs after ten years is shown in Figure 4, B. Over this period, the process is predicted to yield 352.16 kg of biomass and 371.12 kg of oxygen. The highest costs are those attributed to water mining (130.55 kg), followed by those pertaining to structural mass (68.98 kg), thermal control (69.09 kg), crew time (47 kg), power (41.17 kg), carbon dioxide (27.41 kg) and volume (9.16 kg).

Given the many unknowns surrounding a mature cyanobacterium cultivation system, and more broadly long-term, crewed mission to Mars, the Main scenario is hypothetical. The model we developed can be used for predicting productivity under different sets of assumptions. Examples of alternative scenarios are described in Table 1. Their predicted time to BEP, as well the breakdown of their costs and product masses and their specific ESM after ten years, are given in Figure 4, C.

In the “High maintenance” scenario, we assume that reactors face extensive malfunctions and wear, leading to a four-fold increase in crew time and to an increase in mass resulting from the replacement, each year, of parts representing 50% of the reactors’ mass. Among all tested scenarios, this one led to the highest specific ESM after ten years (80% increase from the Main scenario’s) and longest time to BEP.

In the “Biofilm” scenario, cyanobacteria are grown as biofilms rather than planktonic cells. We assume a reduction in water usage, reactor mass, and power/cooling requirements of the pumps by, respectively, 90%, 14% and 92% (*20*). We did not account for physiological differences between biofilms and planktonic cells, or for the constraints biofilms would pose on hardware design. The “Biofilm” scenario reduced the specific ESM after ten years by one third and led to the shortest time to BEP.

In the “Gas ISRU” scenario, carbon and nitrogen are sourced from the Martian atmosphere but nutrients provided from the regolith in the Main scenario are imported from Earth. Product yields are high (54% increase from the Main scenario after ten years) but, due to an over-three-fold increase in costs related to power and thermal control, the time to reach the BEP, as well as the specific ESM after ten years, are slightly higher than these of the Main scenario.

### Impact of changes in the value of selected parameters

In order to assess the impact of changes in the value of selected parameters on specific ESM and biomass production, we performed a sensitivity analysis where values were increased or decreased by 20% from the Main scenario. Results are shown in Figure 5.

**Figure 5.**
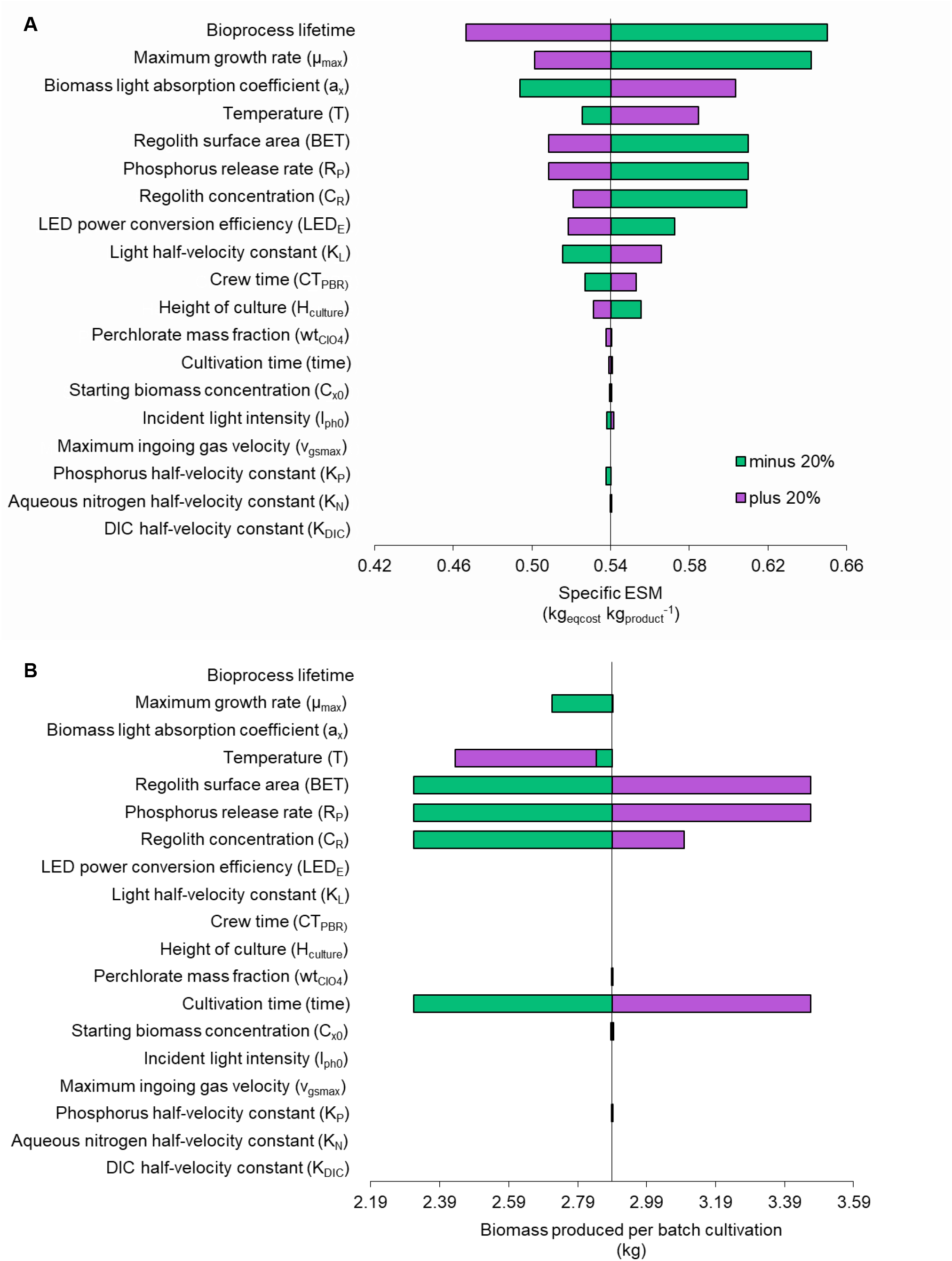
Sensitivity analysis obtained by increasing and decreasing individual input parameters by 20% in the Main scenario. **A**: Effects of input variation on specific ESM over ten years of operation. **B**: Effects of input variation on biomass produced per 30-day batch cultivation run.

The parameter which is most impactful to the specific ESM is the expected operational lifetime of the reactor (“bioprocess lifetime”), due to the high establishment costs and to the subsequent sharp decrease in yearly costs (see Equivalent system mass Section).

Second comes the maximum growth rate. An increase in its value reduces the specific ESM, though biomass production remains unaffected due to phosphorus limitations. On the other hand, a decrease in growth rates leads to a decrease in biomass production. It should however be noted that changes in maximum growth rates only (not accompanied by, e.g., changes in half-velocity constants for light and nutrients) may not accurately reflect the interdependency of this factor with others *in vivo*. Increasing growth rates independently of the half-velocity constants may notably lead to an overestimation of the growth rates permitted by resource availability.

Third is the biomass light absorption coefficient, whose variation affects the ESM_specific_ in the same direction: its decrease leads to a decrease in shading and, therefore, in light intensity requirements at the surface of the reactor. Biomass production is not impacted as phosphorus availability, not light, remains limiting. Changes in the light half-velocity constant have a similar, though less pronounced, effect, and for similar reasons (when it decreases, a given growth rate can be maintained at a lower light intensity). Changes in the power conversion efficiency of LEDs have effects of the same magnitude as changes in the light half-velocity constant, but opposite (with a higher efficiency, less power is required to reach a given light intensity).

A variation in temperature from its optimal value (304.89 K) reduces growth rates, and consequently increases the specific ESM and decreases biomass production.

Variations in phosphorus release rates per surface area, or in regolith surface area, affect biomass productivity and the specific ESM in the same way as they similarly impact phosphorus availability. An increase in either leads to an increase in biomass production and a decrease in specific ESM, and conversely. Changes in regolith concentration affect phosphorus availability in the same fashion, but their effect on biomass production and specific ESM are modulated by the associated change in perchlorate concentration.

Variations in the values of the remaining parameters change the specific ESM by less than 3%.

## DISCUSSION

Bioproduction systems fed through cyanobacteria, themselves grown using resources naturally available on site, could open avenues for sustainable crewed missions to Mars (*12*). Here we provide a mathematical model for predicting the productivity and ESM efficiency of cyanobacterium cultivation through ISRU. Given that key parameters of long-term crewed missions remain to be defined, and that relevant technologies are expected to mature before they take place, this model should be seen as a basis to build upon and the presented predictions, as a snapshot of today’s advancement. Results nonetheless show that the cultivation of cyanobacteria through ISRU can be resource-effective in the long term: we predict a breakeven point within five years in the Main scenario and within ten in all other scenarios. Our model can also help guide further development: relying on it, we underline areas where improvements would most improve resource-efficiency; point out adaptations of existing technology which are required for the foreseen cultivation system to be operated; and list gaps in knowledge whose filling in would enable refined predictions.

Among the main sources of uncertainty is the provision of nutrients from local resources. Most (in terms of diversity) would come from regolith. Here, phosphorus release rates from the MGS-1 regolith analogue were used as a proxy for nutrient availability. This estimate likely errs on the conservative side: the Martian crust is richer in phosphorus than that of the Earth, and the dominant phosphate minerals on Mars are more soluble than their terrestrial counterparts (*30*). That effect will however be limited by the fact that another element may then become limiting (see for instance Figure S1). On the other hand, the regolith on Mars is not perfectly homogeneous. One may rather look for a unit – or a combination of units – with a more suitable grain size and composition. Phosphate-rich materials (*31*) could for instance be used to increase phosphorus contents, or gypsum for sulfur (*16*). Beneficiation could be applied if worth the cost. Overall, the accuracy of our estimates is limited by the scarcity of knowledge on elemental release dynamics from the regolith into water, let alone when cyanobacteria intensify this release (*16, 17*).

While a large variety of elements would come from regolith minerals, circa 97% of the biomass would be composed of elements (C, H, N, O) provided from atmospheric gases and water. Uncertainties could be reduced – and volumetric productivity increased – by bringing from Earth the nutrients which regolith would otherwise provide. A related system (but where nitrogen is imported as well) has been outlined elsewhere (*20*). This approach loses relevance with increasing mission duration but may be more reliable and easier to implement, and thereby more suitable for early missions. Intermediate options are also worth considering. Regolith could for instance be supplemented with imported phosphorus, which only represents about 1.5% of the biomass (Equation 1 in Materials and Methods).

Further uncertainty comes from the provision of gaseous nutrients from the Martian atmosphere. While we here assume that pressurized, nitrogen-enriched gas will be a byproduct of carbon dioxide purification envisioned for, e.g., propellant or polymer production) (*32*), additional processing steps may be required (*14*). On the other hand, dinitrogen would not be the only source of nitrogen: nitrates have been detected in Martian meteorites (*33, 34*) and within Gale crater materials (*35*). As on-site measurements are geographically limited, we omitted nitrates from the present analysis. Including them would, however, only increase the predicted productivity by a small extent: results from measurements on site range from below detection to circa 700 ppm (*36*–*38*). Based on Equation *1* and assuming 300 ppm, at the higher end of estimates for Rocknest samples, nitrogen from nitrates could contribute at most to less than 7% of the biomass produced in in the Main scenario.

In the Main scenario, cyanobacteria are fed exclusively with resources mined on site. One may argue that a cyanobacterium cultivation module beyond Earth would advantageously rely on waste from the habitat, as foreseen within Compartment IV of ESA’s MELiSSA project (*11*). It has also been proposed that, on Mars, urine be used as the main nitrogen source for (in these instances, non-diazotrophic) cyanobacteria (*39, 40*). Here we focused on ISRU since, with recycling alone, amounts of a given element (e.g., nitrogen) can only decrease over time, limiting the autonomy of the system. Besides, other processes such as plant cultivation may compete with cyanobacterium cultivation for waste. We nonetheless expect optimal configurations to include a combination of ISRU and waste processing. As an example, one human releases about five kilograms of nitrogen (four of which in their urine) and 0.75 kg of phosphorus a year (*41*), which corresponds (based on Equation 1) to the nitrogen and phosphorus covering, respectively, 67 and 51 kgs of cyanobacterial biomass – produced in our main scenario within 1.9 and 1.4 years. Recycling could therefore lead to resource-efficiencies which are higher than predicted here.

Knowledge gained through further research can enable more accurate assessments, and technological advance can improve the resource-efficiency of the system we outlined. Directions for both, as identified while developing our model or based on its predictions, are listed in Table 2 – with the hope that they will inspire dedicated research projects. As examples, additional knowledge on the interactions between regolith and cyanobacteria could greatly enhance the accuracy of our predictions; and the development of cultivation systems that minimize water consumption would be instrumental in decreasing resource-efficiency. Our results however suggest that even in the current state of the art, cyanobacterium production from *in situ* resources is a viable foundation for bioproduction on Mars.

**Table 2.** Recommendations to improve the accuracy (A) of future estimates and to improve the resource-efficiency (E) of ISRU-based cultivation of cyanobacteria on Mars.

| To improve estimate accuracy |  |  |
| --- | --- | --- |
| Recommendation |  | Comment |
| A1 | Characterizing the dynamics of nutrient availability in a regolith-cyanobacterium system | Must account for regolith properties (which vary) and cyanobacterium-driven weathering. Greatly affect growth rates, as well as required renewal rates for regolith. Should incorporate progress made on Recommendation A6. |
| A2 | Designing and assessing the mass of a natural light collection and distribution system | Currently poorly defined, though progress made for plant cultivation offers a strong basis. Its mass, and the fraction of light that should come from electric lighting, will largely affect the specific ESM. Should be done in conjunction with E1. |
| A3 | Better defining heating and cooling requirements | Resource-efficiency may be largely impacted by the amount of heat to be evacuated with external systems (e.g., flow-through radiators; here we assumed equivalence to the power consumed), or purposely produced, to maintain the target cultivation temperature. |
| A4 | Refining concepts for long-term, crewed missions to Mars | No detailed reference scenario is available for long-term missions. Key parameters (e.g., ISRU technologies used for other mission components, power sources, needs for consumables) remain undefined. Here we largely relied on technologies considered for shorter, early missions. |
| A5 | Selecting the most relevant downstream applications | Their nature (e.g., production of edible plant biomass, or of bioplastic) will affect the specific ESM of end products, determine recycling needs, and dictate the scale of the cultivation system. Depends on A4. |
| A6 | Further characterizing the physiology of the | In particular, its fitness under factors uncommon on Earth (such as a dependence on relevant types of regolith) and their combination. |
|  | cyanobacterium strain to be used |  |
| A7 | Further characterizing the Martian regolith around potential landing sites | The properties of the regolith to be used (e.g., mineralogy, grain size, toxicity) greatly impact estimates. These properties vary with location. |
| **To improve resource-efficiency** |  |  |
| **Recommendation** |  | **Comment** |
| E1 | Developing a Mars ISRU-specific photobioreactor | Trade-offs differ widely from these to be made on Earth (e.g., costs are largely driven by ESM). Unique constraints are posed by the Martian environment and the need for hauling. Regolith may wear hardware out and shade cells if improperly managed. |
| E2 | Developing efficient systems for the mining of local resources (water, regolith, atmospheric gases) | Given their versatility in crewed missions to Mars, we expect further development and economies of scale which could greatly improve the specific ESM. However, modules (e.g., pertaining to gas processing) dedicated to cyanobacterium cultivation may be needed. |
| E3 | Developing regolith handling systems | Some (e.g., for conveying regolith from extraction sites) will likely be versatile in a long-term Mars mission; others (e.g., transfer of regolith to and from the culture) may be more specific. |
| E4 | Improving automation, robustness and reliability | Increases in requirements for crew time and spare parts would strongly decrease resource-efficiency. |
| E5 | Increasing the efficiency of lighting systems | Artificial lighting generates large costs through power and cooling needs, especially when supporting high growth rates. LED efficiency is expected to increase in the near future; improving light distribution (including of natural light) in the system is another possible approach. LED spectra can be selected for improved power-to-biomass conversion efficiency. |
| E6 | Developing water-efficient hardware and processes | Water represents a large fraction of establishment costs. Biofilms are worth considering. |
| E7 | Developing systems for the downstream processing of cyanobacterium products | Development may be needed, for instance, for purifying biomass when mixed with regolith grains of cell-like size; or for efficiently digesting cyanobacteria for improved used as a nutrient source, notably for plants. Depends on A5. |
| E8 | Reducing hardware mass, volume and power consumption | In conjunction with E1. Numerous instruments have been miniaturized for use beyond Earth, often with dramatic decreases in mass and power use. |
| E19 | Developing regolith beneficiation processes | Nutrient availability from the regolith is the main limiting factor in the Main scenario. |
| E10 | Selecting and engineering cyanobacterium strains for Martian ISRU | Strains more suitable than the one assumed here could be identified. The selected strain could be engineered for, for instance, higher resistance to relevant stress factors or abilities to use local resources. |

Our work also bears relevance to bioproduction on Earth. What is learned and conceived for Mars exploration, where constraints on resources are so extreme that radically novel systems are developed out of necessity, can foster new approaches to sustainability. Here, extensive experiments in realistic environments allowed us to develop a model capable of making quantitative predictions of system performance under unusual constraints and from a unique set of input resources. The same approach could help design systems on Earth which turn raw, abundant resources such as non-arable soils, dinitrogen and carbon dioxide into substrates for bioprocesses, without burning fossil fuels. These may in turn form a foundation for sustainable bioproduction systems, including in resource-poor environments, and help face the environmental challenges of the present century.

## MATERIALS AND METHODS

### Assumed design and operation of the photobioreactor

We assume a typical airlift photobioreactor with a liquid phase of one cubic meter, which contains water, regolith, cyanobacteria, carbon dioxide, dinitrogen, and oxygen (Figure 6). Cultivation starts in batch mode. In this mode, the reactor is a semi-closed system: no liquid flows in or out but carbon dioxide and dinitrogen are continuously gassed. Once a given biomass concentration has been reached, the reactor can switch to continuous mode (turbidostat cultivation). The culture medium is then continuously removed and replenished with a suspension of regolith in water, such that the concentration of biomass, regolith, and volume remain constant. The reactor is lit continuously with sunlight supplemented (e.g., during the night or dust storms) with artificial light.

**Figure 6.**
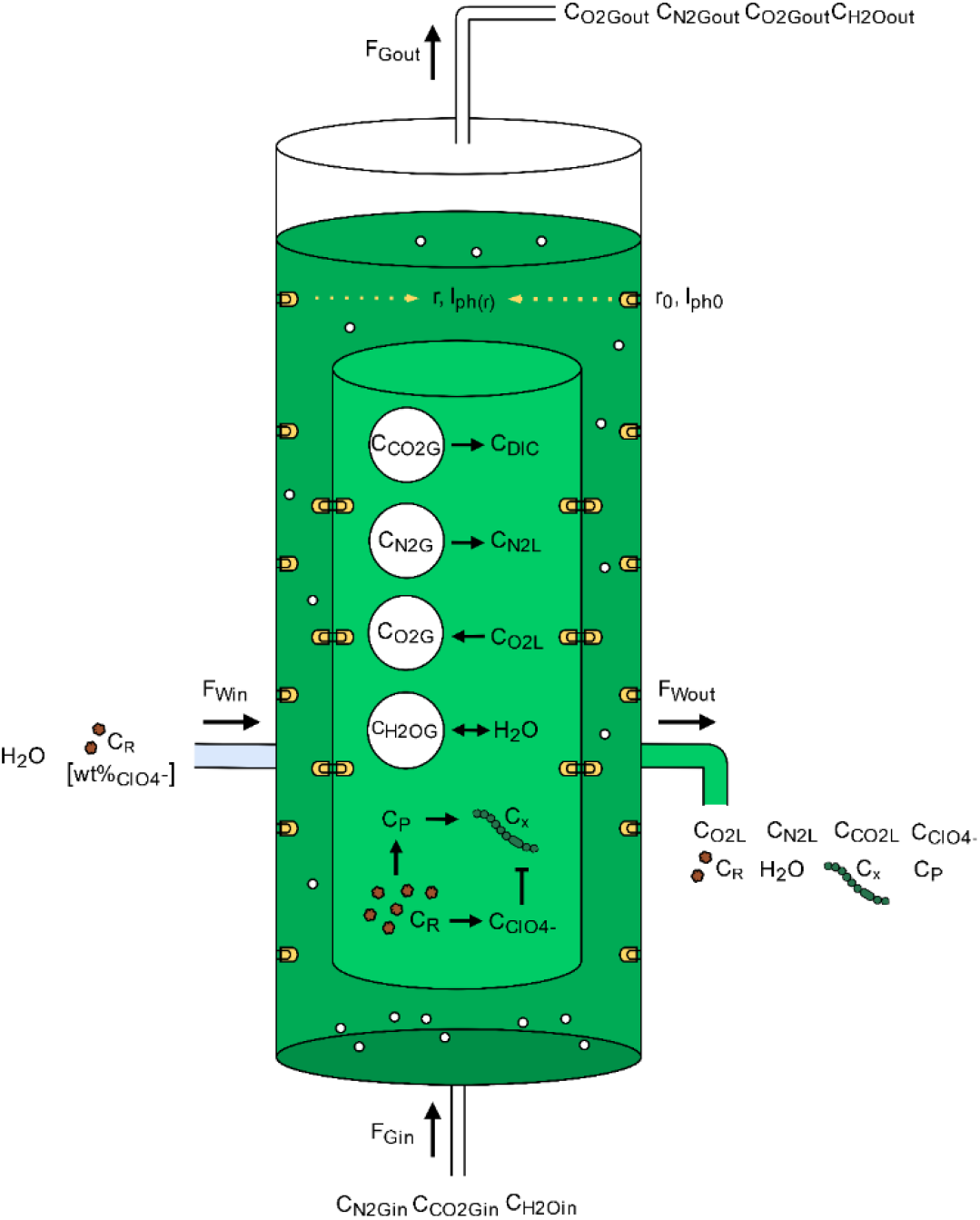
Schematic representation of an airlift photobioreactor for cyanobacterial biomass production from Martian resources. The culture medium consists of water (*H*_*2*_*O*) supplemented with regolith (*C*_*R*_). The regolith releases phosphorus (*C*_*P*_) and perchlorate (*C*_*ClO4-*_), which supports and inhibits, respectively, the growth of cyanobacteria (*C*_*x*_). The gas phase within the photobioreactor contains water (*C*_*H2OG*_), carbon dioxide (*C*_*CO2G*_), nitrogen (*C*_*N2G*_) and oxygen (*C*_*O2G*_). The gas inflow (*F*_*Gin*_) and outflow (*F*_*Gout*_) replenish the dissolved inorganic carbon (*C*_*DIC*_) and dissolved nitrogen (*C*_*N2L*_) required for biomass production, flush out dissolved oxygen (*C*_*O2L*_), and stir the medium. The liquid inflow (*F*_*Win*_) and outflow (*F*_*Wout*_) maintain the biomass concentration constant during operations in continuous mode. Sunlight is provided via light guides and supplemented with artificial light (using LEDs). Light guides and LEDs are placed on the outer shell, radiating inwards, and on the inner shell, radiating in- and out-wards. Light sources have a radial position defined as *r*_*0*_ and a corresponding light intensity (I_ph0_). The light availability decreases with distance from the incident light (*r*), leading to a localized light intensity (*I*_*ph(r)*_).

### Model organism and growth estimation

The model described in the following subsections is based on the cyanobacterium *Anabaena* sp. PCC 7938, which we selected based on its abilities to use resources available on Mars and its suitability as a source of nutrients for downstream bioprocesses (*22*). Parameter values defined for other filamentous cyanobacteria were sourced from literature when not available for this specific strain. We relied, for the modelling of its growth, on the biomass stoichiometric equation for auto- and diazotrophic growth of the closely related *Anabaena cylindrica* PCC 7122, determined by Barbera et al. (*42*) and normalized in Equation 1 by mole of carbon (*mol*_*x*_):

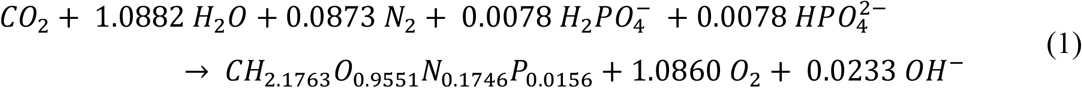

The molar coefficients for oxygen 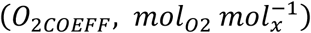, nitrogen 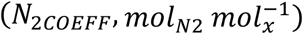 and phosphorus 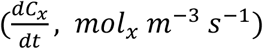 are taken from Equation 1. We complemented this stochiometric information with data available for other filamentous cyanobacteria – *Arthrospira platensis* (*43, 44*) and *Anabaena cylindrica* (*45*) – to account for micronutrients (Table S2), leading to an estimated *Anabaena* sp. PCC 7938 mass per carbon mole (*M*_*x*_) of 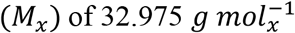.

Biomass productivity 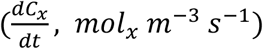 is estimated as the product of biomass concentration (*C*_*x*_, *mol*_*x*_ *m*^−3^) and specific growth rate:

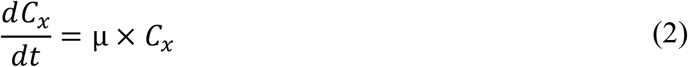

The specific growth rate (μ, *s*^−1^) is calculated as a function of system temperature (*T, K*), light availability (*I*_*ph*_, μ*mol*_*ph*_ *m*^−2^ *s*^−1^), phosphorus concentration (*C*_*P*_, *mol*_*P*_ *m*^−3^), and concentration of perchlorate 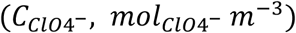:

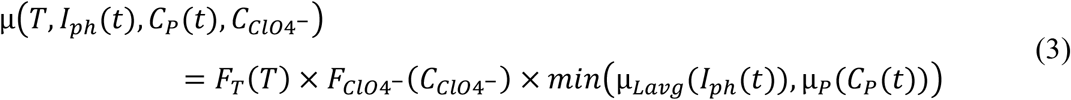

Threshold kinetics are assumed for light and phosphorus: growth at a given time is constrained only by the one among both which is limiting at that time. Consequently, the rightmost term in Equation 3 corresponds to the minimum (MATLAB’s min function) between the growth rate predicted as a function of light intensity (μ_*Lavg*_, *s*^−1^) and that predicted based on phosphorus concentration (μ_*P*_, *s*^−1^). The effects on growth rates of temperature and perchlorate concentration, which are driven by toxicity(*17*) and thermodynamics (*46*), follow multiplicative kinetics and reduce growth by a factor independent of the value of other variables.

### Overview of the model

We developed a photobioreactor-centred model in MATLAB (ver. R2022a), whose structure is depicted in Figure 7. A modular approach was chosen to create an adaptable tool that can be integrated into other code, extended, or otherwise improved. The model is composed of seven modules: Temperature; Production; Regolith and perchlorate; Light 1; Light 2; Gas; and ESM. At the centre of the model is the Production module, which integrates the equations for biomass productivity and specific growth rate (Equations 2 and 3). Other modules are used to generate input for the growth rate calculation (Temperature, Regolith and perchlorate, Light 1), or to estimate the resource requirements (Light 2, Gas, ESM) of biomass and oxygen production. An overview of the computing process is given below.

**Figure 7.**
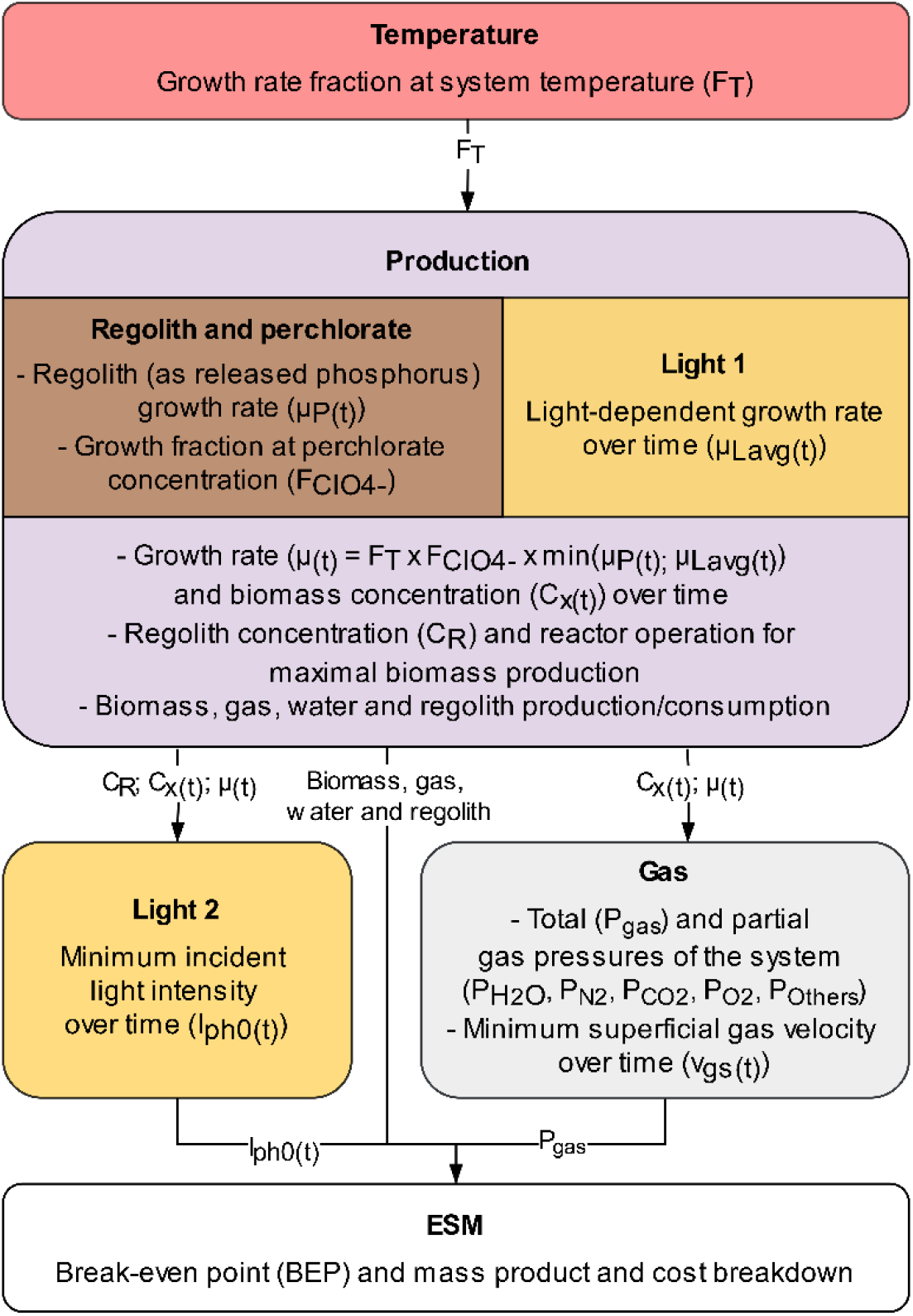
Graphical representation of the model’s structure. Each box represents a module. Arrows indicate variables transferred between modules.

Computing starts with the Temperature module, which determines the fraction of growth rate at a set temperature. This fraction (*F*_*T*_) is fed into the Production module, which includes the Regolith and perchlorate, as well as Light 1, modules. These two modules calculate the growth rates which can be obtained over time as a function of the concentration of regolith – accounting for its being a source of both perchlorates (*F*_*ClO4-*_) and leached phosphorus (*μ*_*P(t)*_) – and of the maximum light intensity at the surface of the reactor (*μ*_*Lavg(t)*_). Both light and regolith-based growth rates are used to determine the optimum regolith concentration (*C*_*R*_) and length of batch mode (before switching to continuous mode) in a run. Relying on these optimum values, the Production module sets the input parameters (growth rate (*μ*_*(t)*_), biomass concentration (*C*_*x*_), and total and time-dependent flows of gas, water, and regolith) for the Light 2 and Gas modules. These in turn determine the light intensity (*I*_*ph0(t)*_), partial pressures of nitrogen (*P*_*N2*_) and carbon dioxide (*P*_*CO2*_), and ingoing gas superficial velocity (*v*_*gs(t)*_), required at a given time to sustain the predicted productivity, as well as the partial pressures of water (*P*_*H2O*_) and oxygen and the total pressure (*P*_*gas*_). Finally, the ESM module computes, based on inputs from all other modules, values pertaining to resource-efficiency. The modules are detailed in the following subsections.

### Temperature module

The temperature-dependent fraction of the maximum growth rate (*F*_*T*_) is calculated according to the model of Bernard and Rémond (*46*):

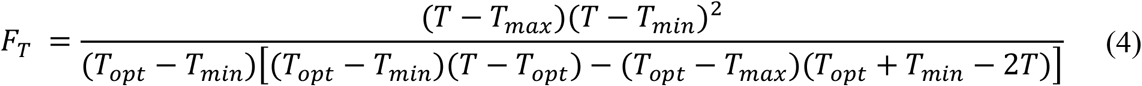

where *T, T*_*max*_, *T*_*opt*_ and *T*_*min*_ correspond to system temperature, maximal growth temperature, optimal growth temperature and minimal growth temperature, respectively. All temperatures are given in Kelvin. We used the maximal, optimal and minimal growth temperature values determined experimentally for *Anabaena sp*. PCC 7122 (*42*).

### Regolith and perchlorate module

Regolith concentrations impact growth dynamics through the interplay of two components: increasing the concentration of regolith increases nutrient availability, but also the concentration of perchlorates in the liquid phase (*17*). The effects of both are determined separately.

### Phosphorus-dependent growth rate

Here, the growth rate is calculated as a function of phosphorus concentration, the nutrient assumed to be limiting in regolith-dependent growth (*17*). Its variation over time 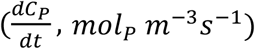 is dictated by the release, consumption and (in continuous mode) dilution rates of phosphorus (Equation 5).

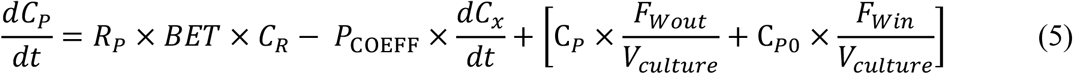

The release term corresponds to the leaching of phosphate from the regolith and is the product of the linear release rate of phosphorus (*R*_*p*_, *mol*_*p*_ *m*^−2^ *s*^−1^), the regolith surface area 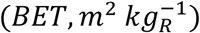 and the regolith concentration (*C*_*R*_, *kg*_*R*_ *m*^−3^)(*45*). Linear release rates of phosphorus were deduced from the amount of biomass produced by *Anabaena* sp. PCC7938 when grown in water supplemented with MGS-1 simulant (Table S3). The regolith surface area used in our Main scenario is that we determined for the MGS-1 simulant (see Supplementary data S6). The consumption term is the biomass productivity 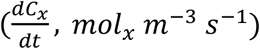 multiplied by the molar coefficient for phosphorus 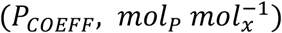. The dilution term (in square brackets) corresponds to the product of phosphorus concentration (*C*_*P*_, *mol*_*P*_ *m*^−3^) and flow of outgoing water (*F*_*Wout*_, *m*^3^ *s*^−1^) per culture volume (*V*_*culture*_, *m*^3^). The replenished water (*F*_*Win*_, *m*^3^ *s*^−1^) per culture volume (*V*_*culture*_, *m*^3^) is also taken into account, although the factor corresponding to its phosphorus concentration (*C*_*P*0_, *mol*_*P*_ *m*^−3^) is assumed to be zero. A phosphorus-dependent, specific growth rate (μ_*p*_, *s*^−1^) is then calculated based on Haldane kinetics (*47*):

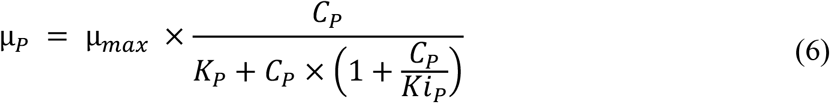

*K*_*P*_ is the phosphorus half velocity constant – the non-inhibitory phosphorus concentration supporting half the maximum growth rate (μ_*max*_, *s*^−1^) – and *K*_*iP*_ the phosphorus inhibition constant: the phosphorus concentration at which the growth rate is decreased, through inhibition, to half of its maximum value. Both constants were previously determined experimentally (*17*).

### Growth rate inhibition by perchlorates

Perchlorate concentration 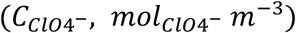 is calculated from the regolith concentration (*C*_*R*_, *kg*_*R*_ *m*^−3^) and perchlorate mass fraction 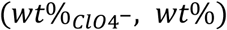. Given the high solubility of perchlorate salts, they are assumed to dissolve promptly into the liquid phase. The fraction of the maximum growth rate obtained at a given perchlorate concentration 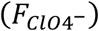 is calculated with the empirical Equation 7 (determined previously) (*17*):

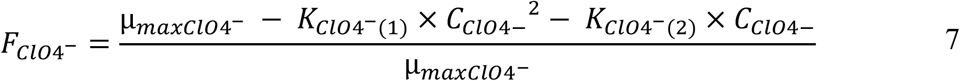

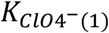 and 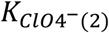 are perchlorate inhibition constants, and 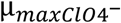 the maximum growth rate documented in the study where they were experimentally determined (*17*).

### Light 1 module

The Light 1 module determines the highest growth rates which can be obtained as function of the average light intensity throughout the culture medium (μ_*Lavg*_, *s*^−1^).

To that end, light availability throughout the photobioreactor is first estimated using Equation 8, based on the Lambert-Beer law for cylindrical bioreactors (*48*):

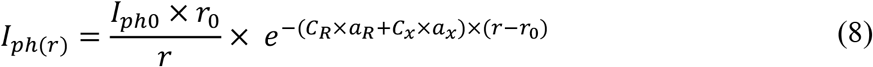

Here, the light intensity at a given radius (*I*_*ph*(*r*)_, *mol*_*ph*_ *m*^−2^ *s*^−1^) is calculated from the incident light intensity (*I*_*ph*0_, *mol*_*ph*_ *m*^−2^ *s*^−1^) as a function of the distance from the light source and of light attenuation. The distance is obtained by subtracting the radial position of the light source (*r*_0_, *m*) from the given radius (*r, m*). The light attenuation is estimated from the spectrally averaged specific light absorption coefficients of biomass 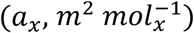 and – if shading by suspended regolith is included – of regolith (*a*_*R*_, *m*^2^ *kg*^−1^), multiplied by their respective concentrations. These coefficients were estimated based on the concentration-dependent attenuation of light intensity by cyanobacteria (Figure S2) or MGS-1 regolith (*17*). In the present study, regolith shading was only considered in the preparation of Figure 1. The weighted average light intensity in the photobioreactor is used to calculate μ_*Lavg*_ based on Monod kinetics (*49*), input constants for which were determined empirically (Figure S3).

### Production module

The Production module starts by determining the optimal regolith concentration and length of the batch (as opposed to continuous) phase which maximizes the overall biomass production. This is achieved using two MATLAB “for loop” functions: an outer loop which covers a range of regolith concentrations, and an inner loop which covers different lengths of batch phase for each regolith concentration.

The outer loop assumes a batch phase for the entire cultivation cycle. Biomass and phosphorus concentrations over time are estimated with MATLAB’s ODE45 function (solving Equations 2, 3, 6 and 8, and light-dependent Monod kinetics provided in Section S6). At each time step of the outer loop, the inner loop switches to continuous mode for the rest of the cultivation cycle. In that phase, the biomass concentration calculated by the outer loop becomes the set biomass concentration (*C*_*xset*_, *mol m* ^−3^), while phosphorus concentration (initially, that calculated by the outer loop) and reactor dilutions are time-dependent variables calculated with an ODE45. The total biomass produced (*Biomass*_*produced*_, *mol*_*x*_) is calculated as the set biomass concentration multiplied by the sum of the culture volume and of the extracted water volume (−*w*_*out*_, *m*^3^):

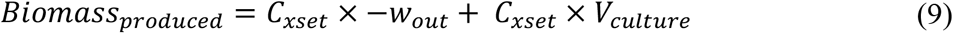

The combination of regolith concentration, and lengths of batch and continuous phases, is recorded. Biomass concentration and growth rates over time under these parameters, as well as the flow rates and total volumes of gas, water and regolith, serve as inputs for the Light 2, Gas and ESM modules.

### Light 2 module

When light is non-limiting, its intensity can be decreased to reduce energy expenditure. The minimum intensity required to sustain the productivity calculated in the Production module is determined within the Light 2 module, using the same set of equations as in the Light 1 module.

### Gas module

The photobioreactor is continuously gassed with a water-saturated mixture of carbon dioxide and dinitrogen. The gas stream provides the carbon and nitrogen required for biomass production, removes oxygen and stirs the medium. Gas-related calculations are divided into two sections. The first section calculates the minimal pressure of each gaseous component needed to maintain the growth rates calculated in the Productivity module. The second section estimates the velocity of the ingoing gas stream necessary to replenish the consumed carbon and nitrogen. To avoid unnecessary complexity, we here assume a bubble column, rather than an airlift, photobioreactor. Although the latter tends to be slightly less efficient in gas-liquid mass transfer (*50*), requirements on gas partial pressures would only be slightly higher and the difference has a negligible effect on resource-efficiency.

The main outputs of this module’s first section are the total (*P*_*gas*_) and partial pressures of the gaseous components (*P*_*N*2_, *P*_*CO*2_, *P*_*O*2_, *P*_*H*2*O*_). All gas pressures are expressed in millibar. The partial pressures of carbon dioxide (which speciates into dissolved inorganic carbon) and dinitrogen correspond to the minimum that ensures neither carbon nor nitrogen are limiting nutrients. To determine them, the corresponding dissolved inorganic carbon (*C*_*DIC*_, *mol*_*DIC*_ *m*^−3^) and dissolved nitrogen concentrations (*C*_*N*2*L*_, *mol*_*N*2*L*_ *m*^−3^) are first calculated assuming Monod kinetics (Equations 10 and 11):

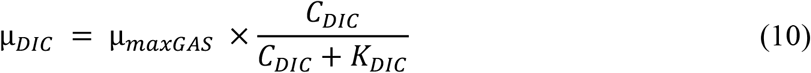

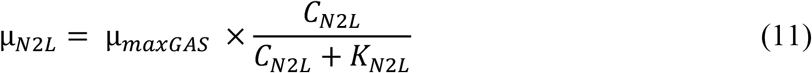

The half velocity constants for dissolved inorganic carbon (*K*_*DIC*_, *mol*_*DIC*_ *m*^−3^) and dissolved nitrogen (*K*_*N*2*L*_, *mol*_*N*2*L*_ *m*^−3^), as well as the associated maximum growth rate (μ_*maxGAS*_, *s*^−1^), were determined previously (*14*). The growth rates dictated by the concentrations of dissolved inorganic carbon (μ_*DIC*_, *s*^−1^) or dissolved nitrogen (μ_*N*2*L*_, *s*^−1^) are defined as equal to the growth rates predicted by the Production module.

Maintaining dissolved inorganic carbon and dissolved nitrogen concentrations at, or above, the calculated value requires replenishing these gases as they are consumed. Conversely, dissolved dioxygen (*C*_*O*2*L*_, *mol*_*O*2*L*_ *m*^−3^) must be continuously removed. To determine the necessary gas transfer rates (*iTR, mol*_*iL*_ *m*^−3^*s*^−1^), a mass balance is set up for the liquid phase (Equation 12A). There, *i* stands for *CO*_2_, *N*_2_ or *O*_2_.

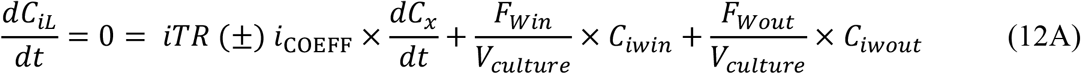

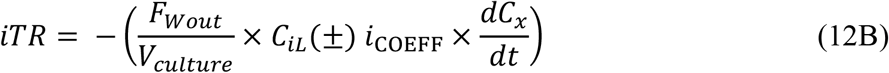

As shown in Equation 12A, four elements determine the evolution over time of the dissolved concentration of each gas: gas transfer from gas to liquid phase (*iTR, mol*_*iL*_ *m*^−3^ *s*^−1^); gas consumption (CO_2_, N_2_) or release (O_2_) by the cyanobacteria, calculated as the product of biomass productivity 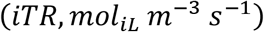 and of the gas’s molar coefficient 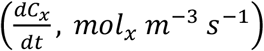 and the dissolved gas entering and exiting with liquid flows, which correspond to the product of the flow (*F*_*w*_, *m*^3^ *s*^−1^) and of the gas concentration therein (*C*_*iw*_, *mol*_*i*_ *m*^−3^) corrected for culture volume. Dissolved concentrations remain constant, hence 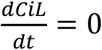. Dissolved gas concentrations in the outgoing flow are assumed to be equal to these inside the reactor (*C*_*iwout*_ = *C*_*iL*_), and the ingoing water is assumed to be pure (*C*_*iwin*_ = 0). Gas-liquid transfer rates are thus calculated according to Equation 12B.

The gas transfer rates (*iTR*) are used to calculate the gaseous concentrations of O_2_, N_2_ and CO_2_ at the liquid column centre (*C*_*iG*_, *mol*_*i*_ *m*^3^):

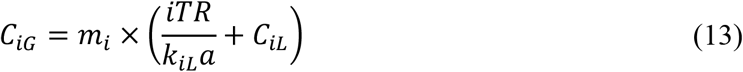

The gas-liquid mass transfer (*ki*_*L*_*a*) under Martian gravity and at under the inner photobioreactor pressure are determined as previously described (*14*). The partition coefficient 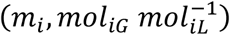 for each gas is calculated using Henry’s law (using the dissolved inorganic carbon and dissolved nitrogen concentrations calculated as described above). For the dissolved oxygen concentration, we rely on the smallest value that results in the computation of a positive oxygen gaseous concentration. The internal partial pressure of each gas is obtained from its concentration with the ideal gas law.

Alongside O_2_, N_2_ and CO_2_, water vapour contributes to the total pressure inside the reactor. Its partial pressure is calculated with standard methodology (*51*–*53*), as previously described (*14*). The total gas pressure inside the reactor is approximated to the sum of the partial pressures (Dalton’s law):

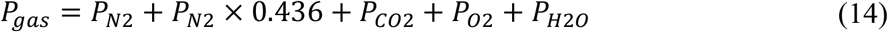

The term *P*_*N*2_ × 0.436 accounts for gases introduced together with nitrogen as impurities. The composition of this gas mixture (56.42% N_2_, 42.07% Ar, and 1.51% CO) corresponds to Martian atmosphere from which carbon dioxide and oxygen were removed (*54*) (carbon dioxide and oxygen purification is foreseen on Mars for other ISRU processes). These impurities are included as we assume no further purification of nitrogen.

In order to minimize the costs associated with the provision of gases, the ingoing superficial gas velocity (v_gs_) should be kept to the minimum required for sustaining the productivity predicted by the Production module. This velocity is calculated as previously described (*14*), using as inputs the total pressures calculated above and the oxygen productivity from the Production module.

### Equivalent system mass module

The resource-efficiency of cyanobacterium production through ISRU is evaluated on the basis of equivalent system mass (ESM). For this, the mass (*M, kg*), volume (*V, m*^3^), power requirements (*P, kW*), cooling requirements (*C, kW*_*th*_), and crew time (*CT*_*pBR*_, *CM*_*h*_) of the system are assessed and converted to mass through mass equivalency factors for volume (*V*_*eq*_, *kg m*^−3^), power requirements (*P*_*eq*_, *kg kW*^−1^), cooling requirements 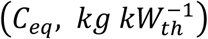, and crew time 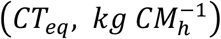. The ESM of the photobioreactor, excluding consumables, is calculated as follows:

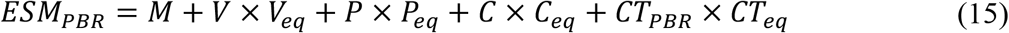

How mass and power requirements are assessed is detailed in Section S8. Cooling requirements are assumed to match power consumption (*55*). In our Main scenario, the volume per photobioreactor is one cubic meter and the required crew time is five hours per year per photobioreactor, assuming a high degree of automation and control from Earth (for comparison: an optimized plant production chamber has been estimated to require 13 hours per square meter per year (*56*)).

Consumables included in the analysis are water, regolith, carbon dioxide and dinitrogen. Water-associated costs are estimated as detailed in Section S8. They are treated as establishment costs since most of the water must be provided at onset and we assume a recycling efficiency close to 100%, an extent achieved with more complex systems and with more stringent constraints on water quality (safety for human consumption)(*57*). The costs of the remaining consumables (*ESM*_*consumables*_, *kg*) are calculated separately:

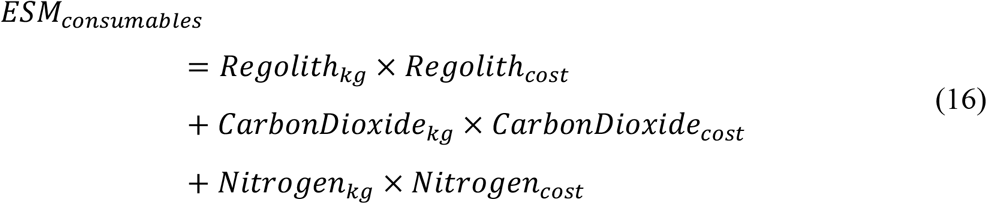

In this equation, the required mass of consumables is multiplied by their respective cost per mass. Information on the determination of this cost is provided in Section S8.

The specific ESM value 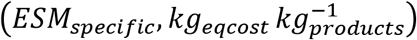 for the bioprocess is:

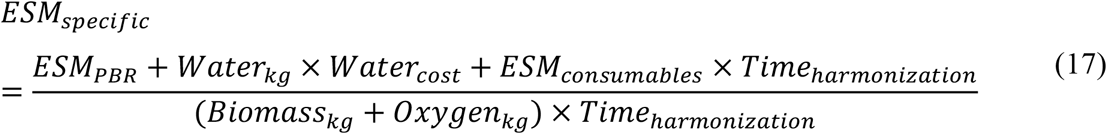

A time correction factor (*Time*_*harmonization*_) is applied to terms corresponding to consumables, biomass, and oxygen, to adjust values computed over a 30-day cultivation period to the entire bioprocess lifetime.

## Acknowledgments

We express our gratitude to Prof. Marcel Janssen (AlgaePARC, Wageningen University and Research) for his support during the early stages of model development. We thank Dr. Depanjalee Dutta and Dr. Michael Maas (Advanced Ceramics group, University of Bremen) for conducting the BET measurements.

## Author contributions

Conceptualization: TPR, CV; Investigation: TPR, DR, EB, BL, CV; Methodology: TPR, CV; Software development: TPR, VB, NK, CH, MA; Writing – original draft: TPR, CV; Writing – review & editing: TPR, VB, NK, DR, EB, BL, GP, CH, SK, MA, CV.

## Data and materials availability

Any data that support the findings of this study will be made available by the authors upon reasonable request. The MATLAB code for the model developed as part of this study was uploaded to GitHub (https://github.com/TPRamalho/BioISRU-Models.git). Enquiries regarding its utilization are welcomed by the authors.

