## Supplementary Materials for "Resource-efficiency of cyanobacterium production on Mars: Assessment and paths forward"

Tiago P. Ramalho *et al.*

**This PDF file includes:**

Supplementary Text in sections S2 to S8
Figs. S1 to S3
Tables S1 to S3
References (58 to 65)

### S1. List of symbols

**Table S1. List of symbols employed in the manuscript with their corresponding description, value (if applicable), unit and reference.**

| Symbol | Description | Value | Unit | Reference |
|---|---|---|---|---|
| $\mu$ | Specific growth rate | | $s^{-1}$ | |
| $\mu_{DIC}$ | Specific growth rate at a given dissolved inorganic carbon concentration | | $s^{-1}$ | |
| $\mu_{Lavg}$ | Average specific growth rate in the photobioreactor as a function of available light | | $s^{-1}$ | |
| $\mu_{max}$ | Maximum growth rate | $8.912 \times 10^{-6}$ | $s^{-1}$ | See Section S6 |
| $\mu_{maxClO4-}$ | Maximum growth rate from cited perchlorate dynamics experiment | $5.042 \times 10^{-6}$ | $s^{-1}$ | [17] |
| $\mu_{maxGas}$ | Maximum growth rate from cited gas utilization experiments | $1.22 \times 10^{-5}$ | $s^{-1}$ | [14] |
| $\mu_{N2L}$ | Specific growth rate at a given dissolved nitrogen concentration | | $s^{-1}$ | |
| $\mu_P$ | Specific growth rate at a given phosphorus concentration | | $s^{-1}$ | |
| $a_r$ | Regolith spectrally averaged specific light absorption coefficient | 15.836 | $m^2\ kg^{-1}$ | [17] |
| $a_x$ | Biomass spectrally averaged specific light absorption coefficient | 3.3145 | $m^2\ mol_x^{-1}$ | This study |
| BET | Regolith surface area | 7605.4 | $m^2\ kg^{-1}$ | This study |
| $Biomass_{kg}$ | Total biomass produced in kg | | kg | |
| $Biomass_{produced}$ | Total biomass produced in moles | | $mol_x$ | |
| C | System cooling requirements | $kW_{th}$ | | |
| $CarbonDioxide_{cost}$ | ISRU Carbon dioxide-to-mass conversion factor | 0.0583 | $kg\ kg_{CO2}^{-1}$ | See Section S8.3 |
| $CarbonDioxide_{kg}$ | Total carbon dioxide input into the photobioreactor | | kg | |
| $C_{ClO4^-}$ | Perchlorate concentration | | $mol\ m^{-3}$ | |
| $C_{CO2G}$ | Gaseous concentration of carbon dioxide in the reactor | | $mol_{CO2}\ m^{-3}$ | |
| $C_{DIC}$ | Concentration of dissolved inorganic carbon | | $mol_{DIC}\ m^{-3}$ | |
| $C_{eq}$ | Cooling power-to-mass conversion factor | 146 | $kg\ kWth^{-1}$ | [56] |

| | | | | |
|---|---|---|---|---|
| $C_{iG}$ | Gaseous concentration of element i | | $mol_i\ m^{-3}$ | |
| $C_{iGin}$ | Ingoing gaseous concentration of element i | | $mol_i\ m^{-3}$ | |
| $C_{iGout}$ | Outgoing gaseous concentration of element i | | $mol_i\ m^{-3}$ | |
| $C_{iL}$ | Dissolved concentration of element i | | $mol_i\ m^{-3}$ | |
| $C_{iwin}$ | Dissolved concentration of element i in the ingoing water flow | 0 | $mol_i\ m^{-3}$ | |
| $C_{iwout}$ | Dissolved concentration of element i in the outgoing water flow | | $mol_i\ m^{-3}$ | |
| $C_{N2L}$ | Dissolved concentration of nitrogen | | $mol_{N2}\ m^{-3}$ | |
| $C_{N2G}$ | Gaseous concentration of nitrogen in the reactor | | $mol_{N2}\ m^{-3}$ | |
| $C_{O2L}$ | Dissolved concentration of oxygen | | $mol_{O2}\ m^{-3}$ | |
| $C_{O2G}$ | Gaseous concentration of oxygen in the reactor | | $mol_{O2}\ m^{-3}$ | |
| $C_P$ | Phosphorus concentration | | $mol_P\ m^{-3}$ | |
| $C_{P0}$ | Initial phosphorus concentration | 0 | $mol_P\ m^{-3}$ | |
| $C_R$ | Regolith concentration | | $kg\ m^{-3}$ | |
| $CT_{PBR}$ | Crew member hours per year | 5 | $CMh^{-1}$ | |
| $CT_{eq}$ | Crew member hours-to-mass conversion factor | 0.94 | $kg\ CMh^{-1}$ | 56 |
| $C_x$ | Biomass concentration | | $mol_x\ m^{-3}$ | |
| $C_{x0}$ | Initial biomass concentration | 0.6214 | $mol_x\ m^{-3}$ | |
| $C_{xset}$ | Biomass concentration during the production phase | | $mol_x\ m^{-3}$ | |
| $ESM_{consumables}$ | Cost of consumables to operate the photobioreactor (regolith, nitrogen and carbon dioxide) | | kg | |
| $ESM_{PBR}$ | Mass cost of implementing and operating (in terms of structure maintenance, power, cooling and crew time) the photobioreactor | | kg | |
| $ESM_{specific}$ | Mass ratio between cost-equivalent and products | | $kg_{eqcost}\ kg_{product}^{-1}$ | |
| $F_{ClO4^-}$ | Fraction of maximum growth rate maintained with a given perchlorate concentration | | | |

| | | | | |
|---|---|---|---|---|
| $F_{Gin}$ | Flow of gas going into the reactor | | $m^3\ s^{-1}$ | |
| $F_{Gout}$ | Flow of gas going out of the reactor | | $m^3\ s^{-1}$ | |
| $F_T$ | Fraction of maximum growth rate maintained at a given temperature | | | |
| $F_{Win}$ | Water flow into the reactor | | $m^3\ s^{-1}$ | |
| $F_{Wout}$ | Water flow out of the reactor | | $m^3\ s^{-1}$ | |
| $H_{culture}$ | Height of culture | 1.7 | m | |
| $i_{COEFF}$ | Biomass molar content of nutrient i per mole biomass | | $mol_i\ mol_x^{-1}$ | |
| $I_{ph0}$ | Light intensity at location r = 0 i.e. incident light intensity | | $mol_{ph}\ m^{-2}\ s^{-1}$ | |
| $I_{ph(r)}$ | Light intensity at a given radius r | | $mol_{ph}\ m^{-2}\ s^{-1}$ | |
| iTR | Gas-liquid transfer rates of element i | | $mol_{iL}\ m^{-3}\ s^{-1}$ | |
| $K_{ClO4^-(1)}$ | Perchlorate inhibition constant 1 | $1.124 \times 10^{-8}$ | | 17 |
| $K_{ClO4^-(2)}$ | Perchlorate inhibition constant 2 | $5.703 \times 10^{-8}$ | | |
| $K_{DIC}$ | Dissolved inorganic carbon half-velocity constant | 0.0457 | $mol_{DIC}\ m^{-3}$ | 14 |
| $k_{iL}a$ | Element i gas-liquid mass transfer coefficient | | $s^{-1}$ | |
| $Ki_P$ | Phosphorus inhibition constant | 1.379 | $mol_P\ m^{-3}$ | 17 |
| $K_L$ | Light half-velocity constant | $2.498 \times 10^{-5}$ | $mol_{ph}\ m^{-2}\ s^{-1}$ | This study |
| $K_{N2L}$ | Nitrogen half-velocity constant | 0.0326 | $mol_{N2}\ m^{-3}$ | 14 |
| $K_P$ | Phosphorus half-velocity constant | 0.0149 | $mol_P\ m^{-3}$ | 17 |
| $LED_E$ | LED light intensity to Joule | $1.66\times10^{-6}$ | $mol\ J^{-1}$ | 64 |
| Bioprocess lifetime | Expected bioprocess operation time | 10 | years | |
| M | Mass | | kg | |
| $m_i$ | Element i partition coefficient gas over liquid | | $mol_{iG}\ mol_{iL}^{-1}$ | |
| $M_x$ | Mass per C-mol biomass | 32.975 | $g\ mol^{-1}$ | This study |
| $N_{2COEFF}$ | Mole nitrogen consumed per C-mol biomass | 0.0873 | $mol_{N2}\ mol_x^{-1}$ | 42 |
| $Nitrogen_{cost}$ | ISRU Nitrogen-to-mass conversion factor | 0 | $kg\ kg_{N2}^{-1}$ | See Section S8.3 |
| $Nitrogen_{kg}$ | Total nitrogen input into the photobioreactor | | kg | |

| | | | | |
|---|---|---|---|---|
| $O_{2COEFF}$ | Mole oxygen produced per C-mol biomass | 1.0860 | $mol_{O2}\ mol_x^{-1}$ | [42] |
| $Oxygen_{kg}$ | Total oxygen produced by the photobioreactor | | kg | |
| P | System power requirements | | $kW^{-1}$ | |
| $P_{CO2}$ | Gaseous pressure of carbon dioxide in the reactor | | mbar | |
| $P_{COEFF}$ | Biomass phosphorus to carbon molar ratio | 0.0156 | $mol_P\ mol_x^{-1}$ | [59] |
| $P_{eq}$ | Power-to-mass conversion factor | 87 | $kg\ kWe^{-1}$ | [56] |
| $P_{gas}$ | Pressure of gas excluding hydrostatic pressure | | mbar | |
| $P_{H2O}$ | Water vapour pressure | | mbar | |
| $P_{N2}$ | Gaseous pressure of nitrogen in the reactor | | mbar | |
| $P_{O2}$ | Gaseous pressure of oxygen in the reactor | | mbar | |
| $P_{Others}$ | Gaseous pressure of gases introduced with nitrogen | | mbar | |
| r | Radius for a given reactor position r relative to the light source | | m | |
| $r_0$ | Radius for the position of light source | | m | |
| $Regolith_{cost}$ | ISRU Regolith-to-mass conversion factor | 0 | $kg\ kg_{Regolith}^{-1}$ | See Section S8.3 |
| $Regolith_{kg}$ | Total regolith input into the photobioreactor | | kg | |
| $R_P$ | Phosphorus release rate | $3.442 \times 10^{-13}$ | $mol_P\ m^{-2}\ s^{-1}$ | See Section S4 |
| T | System temperature | 304.89 | K | |
| time | Cultivation cycle | 2592000 (30 days) | s | |
| $Time_{harmonization}$ | Correction for different timeframes between previous modules and ESM module | | | |
| $T_{min}$ | Minimal growth temperature | 278.44 | K | |
| $T_{max}$ | Maximal growth temperature | 316.08 | K | [42] |
| $T_{opt}$ | Optimal growth temperature | 304.89 | K | |
| V | Volume | | $m^3$ | |
| $V_{culture}$ | Culture volume | 1 | $m^3$ | |
| $V_{eq}$ | Volume-to-mass conversion factor | 9.16 | $kg\ m^{-3}$ | [56] |

| | | | | |
|---|---|---|---|---|
| $v_{gs}$ | Superficial gas velocity | | $m\ s^{-1}$ | |
| $v_{gsmax}$ | Maximal superficial gas velocity | 0.08 | $m\ s^{-1}$ | |
| $Water_{cost}$ | ISRU water-to-mass conversion factor | 0.1290 | $kg\ kg_{H2O}^{-1}$ | See Section S8.3 |
| $Water_{kg}$ | Initial water input into the reactor | | kg | |
| $-w_{out}$ | Total volume of reactor dilutions | | $m^3$ | |
| $wt\%_{ClO4-}$ | Regolith's perchlorate mass fraction | 0.4 | wt% | |

### S2. Effects of phosphorus supplementation on regolith-dependent growth

As phosphorus is the nutrient which, among those provided by regolith, is most likely to be limiting (at least when relying on the MGS-1 simulant (*17*)), phosphorus supplementation can improve growth. The upper limit to this improvement is determined by the phosphorus concentration above which another nutrient becomes limiting. We assessed the impact of phosphorus supplementation on the growth dynamics of *Anabaena* sp. PCC 7938 when relying on MGS-1 as a nutrient source. Cultivation was performed as previously described (*17*) but under a light intensity of 80 $\mu mol_{ph}\ m^{-2}\ s^{-1}$, with concentrations of MGS-1 (purchased from Exolith Lab, Orlando, Florida; shipped May 2019) of 50 and 200 $g\ L^{-1}$ and with supplementation of phosphorus at concentrations ranging from 0 to 0.260 $mol\ m^{-3}$. Biomass productivity ($x_r, \mu g\ m^{-3}\ s^{-1}$) can be decomposed as:

$$x_r = x_r' + x_{r,init} \qquad \text{(SE1)}$$

where $x_r$ and $x_{r,init}$ ($\mu g\ m^{-3}\ s^{-1}$) are, respectively, the productivity with and without supplementation and $x_r'$ the increase in productivity attributed to phosphorus supplementation. A modified Monod equation can then be applied:

$$x_r' = \frac{x_{r,max} \times C_P'}{K_P' + C_P'} \qquad \text{(SE2)}$$

where $C_P'$ ($mol\ m^{-3}$) is the concentration of supplemented phosphorus, $K_P'$ ($mol\ m^{-3}$) a half-velocity constant determined empirically, and $x_{r,max}$($\mu g\ m^{-3} s^{-1}$) the productivity under optimal phosphorus supplementation.

Results are shown in Figure S1.

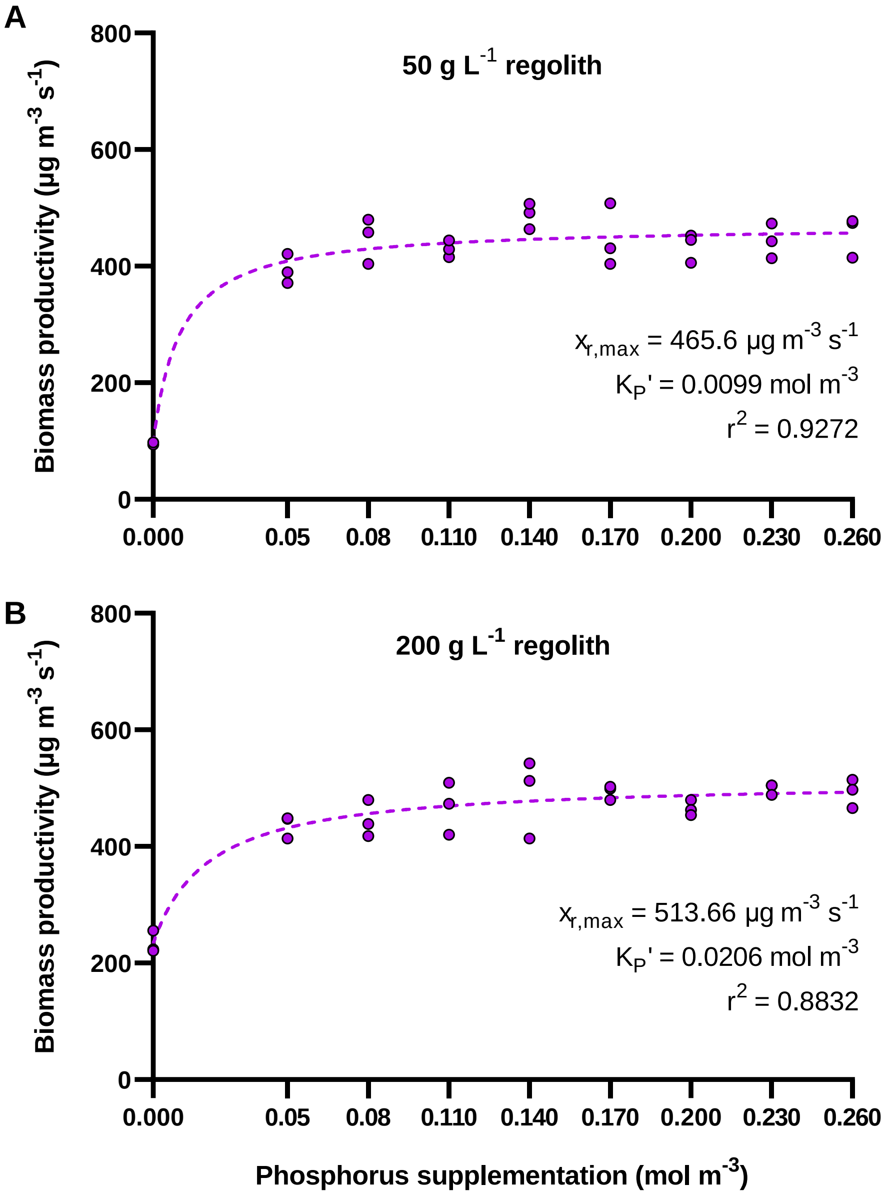


**Figure S1. Growth rates of *Anabaena* sp. PCC 7938 in water containing 50 g $L^{-1}$ (A) or 200 g $L^{-1}$ (B) MGS-1 as a function of supplemented phosphorus.** Dots represent experimental data and the dashed line, the fitted equation, derived from Monod kinetics as described in the text.

**S3. Molar mass per carbon mole ($M_x$)**

**Table S2. Element abundance and mass per carbon mole (C-mol) of *Anabaena* sp. PCC 7938. The molar mass of the biomass (32.965 g $mol_x^{-1}$) is obtained by summing the mass of individual elements.** Input data was sourced from studies pertaining to closely related cyanobacteria (*Anabaena* sp. PCC 7122 and *Anabaena cylindrica* PCC 6309) or other filamentous cyanobacteria (*Arthrospira platensis* PCC 8005).

| Element | Mole of element per C-mol of biomass (mol $mol_x^{-1}$) | Element mass per C-mol of biomass (g $mol_x^{-1}$) | Cyanobacterium | Reference |
|---|---|---|---|---|
| C | 1 | 1.20E+01 | *Anabaena* sp. PCC 7122 | (*42*) |
| H | 2.18 | 2.20E+00 | | |
| O | 9.55E-01 | 1.53E+01 | | |
| N | 1.75E-01 | 2.45E+00 | | |
| P | 1.56E-02 | 4.83E-01 | | |
| S | 7.00E-03 | 2.24E-01 | *Arthrospira platensis* PCC 8005 | (*44*) |
| Fe | 2.80E-04 | 1.56E-02 | | |
| Cu | 1.50E-06 | 9.53E-05 | | (*43*) |
| Mn | 1.40E-05 | 7.69E-04 | | |
| Zn | 3.30E-06 | 2.16E-04 | | |
| K | 7.94E-03 | 3.10E-01 | *Anabaena cylindri ca* PCC 6309 | (*45*) |
| Mg | 1.73E-05 | 4.20E-04 | | |
| Ca | 3.77E-05 | 1.51E-03 | | |

**S4. Phosphorus release rate ($R_P$)**

Linear release rates of phosphorus ($R_P, mol_P\ m^2 s^{-1}$) were estimated from the amount of biomass produced in regolith-based experiments (*17*) (Table S2), using the following equation:

$$R_P = \frac{\left(C_{x(t)} - C_{x0}\right)}{M_{wP} \times (t \times 86400) \times C_R \times BET} \times \frac{g_P}{g_x} \qquad \text{(SE3)}$$

In this equation, the produced biomass $\left(C_{x(t)} - C_{x0}, g\ L^{-1}\right)$ is multiplied by the ratio between mass of phosphorus and biomass ($\frac{g_P}{g_x}, g_P {g_x}^{-1}$) to obtain released phosphorus. The phosphorus is then divided by its molar mass ($M_{wP}, g\ mol^{-1}$), time elapsed ($t \times 86400, s$), regolith concentration ($C_R, kg\ L^{-1}$), and regolith surface area ($BET, m^2\ kg^{-1}$). It is worth noting that various phenomena may occur that limit the accuracy of this approach, such as a decrease in the phosphorus fraction of the biomass or the recapture of released phosphorus through, for instance, the adsorption of phosphate onto iron and aluminum oxides.

**Table S3. Calculation of linear release rates of phosphorus with input data from Ramalho et al. (*17*)**. The phosphorus release rate is calculated per regolith concentration for all timepoints but, for averaged values, only rates documented in periods of vigorous growth (highlighted in green) are considered. The value used as input for the model is the overall average.

Biomass ($g\ L^{-1}$)

| Time (days) | Regolith concentration ($kg\ L^{-1}$) | | | | |
|---|---|---|---|---|---|
| | 0.0125 | 0.025 | 0.050 | 0.100 | 0,200 |
| 0 | 0.063 | 0.063 | 0.027 | 0.027 | 0.027 |
| 3 | 0.069 | 0.071 | 0.105 | 0.090 | 0.087 |
| 7 | 0.126 | 0.161 | 0.126 | 0.122 | 0.110 |
| 10 | 0.194 | 0.217 | 0.235 | 0.238 | 0.231 |
| 14 | 0.227 | 0.268 | 0.296 | 0.346 | 0.380 |
| 21 | 0.213 | 0.268 | 0.528 | 0.566 | 0.646 |
| 28 | 0.177 | 0.302 | 0.532 | 0.714 | 0.775 |

Phosphorus release rate ($mol_P\ m^{-2}\ s^{-1}$)

| Time (days) | Regolith concentration ($kg\ L^{-1}$) | | | | |
|---|---|---|---|---|---|
| | 0.0125 | 0.025 | 0.050 | 0.100 | 0.200 |
| 0 | 0 | 0 | 0 | 0 | 0 |
| 3 | $1.193 \times 10^{-13}$ | $8.868 \times 10^{-14}$ | $4.004 \times 10^{-13}$ | $1.622 \times 10^{-13}$ | $7.665 \times 10^{-14}$ |
| 7 | $5.557 \times 10^{-13}$ | $4.330 \times 10^{-13}$ | $2.176 \times 10^{-13}$ | $1.040 \times 10^{-13}$ | $4.530 \times 10^{-14}$ |
| 10 | $8.107 \times 10^{-13}$ | $4.745 \times 10^{-13}$ | $3.199 \times 10^{-13}$ | $1.626 \times 10^{-13}$ | $7.863 \times 10^{-14}$ |
| 14 | $7.235 \times 10^{-13}$ | $4.516 \times 10^{-13}$ | $2.965 \times 10^{-13}$ | $1.755 \times 10^{-13}$ | $9.710 \times 10^{-14}$ |
| 21 | $4.412 \times 10^{-13}$ | $2.013 \times 10^{-13}$ | $3.678 \times 10^{-13}$ | $1.978 \times 10^{-13}$ | $1.136 \times 10^{-13}$ |
| 28 | $2.514 \times 10^{-13}$ | $2.640 \times 10^{-13}$ | $2.780 \times 10^{-13}$ | $1.891 \times 10^{-13}$ | $1.030 \times 10^{-13}$ |

Average phosphorus release rate ($mol_P\ m^{-2}\ s^{-1}$)

| | | | | | |
|---|---|---|---|---|---|
| Per regolith concentration | $6.966 \times 10^{-13}$ | $4.530 \times 10^{-13}$ | $3.204 \times 10^{-13}$ | $1.652 \times 10^{-13}$ | $8.571 \times 10^{-14}$ |
| Overall | | | $3.442 \times 10^{-13}$ | | |

### S5. Spectrally-averaged biomass light absorption coefficient

The spectrally-averaged biomass light absorption coefficient ($a_x, m^2 mol_x^{-1}$) was obtained as described elsewhere (*59*). In short, three separate *Anabaena* sp. PCC 7938 cultures were washed and resuspended in distilled water. The absorbance of each culture was immediately measured from 400 to 700 nm and at 750 nm. The absorbance value at 750 nm was subtracted from these measurements as it corresponds to scattered light which, unlike absorbed light, can be used by other cyanobacterial cells in the culture. The resulting absorbance value were divided by the light path $(l, m)$ and the biomass concentration $(C_x, mol\, m^{-3})$ to obtain the wavelength-specific biomass light absorption coefficients:

$$a_{x(400-700)} = \frac{(A_{400-700} - A_{750})}{l \times C_x} \qquad \text{(SE4)}$$

Results (including the spectrally-averaged value) are shown in Figure S2.

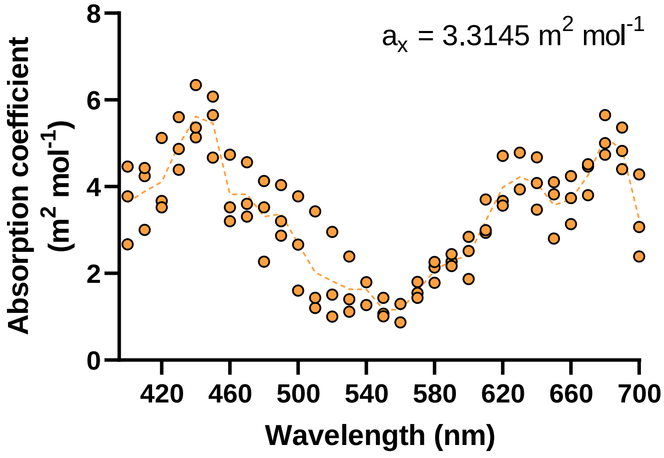


**Figure S2. Absorption coefficient of *Anabaena* sp. PCC 7938 as a function of wavelength.** $a_x$ is the average value over the 400–700 nm range.

### S6. Light-dependent growth kinetics

We determined the dependence on light of *Anabaena* sp. PCC 7938's growth kinetics, as it was critical to the development of the model. *Anabaena* sp. PCC 7938 was grown at 30°C in a Multi-Cultivator MC 1000-OD (Photon Systems Instruments, Drásov, Czech Republic). Each vial of the device was filled with 70 mL of standard, nitrate-free cyanobacterium cultivation medium ($BG11_0$) with 20 mM HEPES (pH 7.1) and flushed continuously with a mixture of $N_2$ (98.5%) and $CO_2$ (1.5%) at a rate of ca. 1 $m_{gas}^3$ $m_{culture}^{-3}$ $min^{-1}$. Warm white light was provided continuously at an intensity of 20, 40, 60, 80, 100, 120, 150 or 170 μ$mol_{ph}$ $m^{-2}$ $s^{-1}$, in triplicate. Cultures were inoculated to an $OD_{750}$ of 0.02–0.05 and sampled daily for 7 days. Growth rates were calculated based on OD evolution during the exponential growth phase. Results (Figure S3) were fitted to commonly-assumed light-dependent growth kinetics: Jassby and Platt (*60*) ($r^2$ = 0.875); Webb (*61*) ($r^2$ = 0.944); and Monod (*49*) ($r^2$ = 0.958). Monod kinetics were assumed for our model:

$$\mu_{Lavg} = \mu_{max} \times \frac{I_{ph}}{K_L + I_{ph}} \qquad \text{(SE5)}$$

In Equation S5, the average light-dependent growth rate $\left(\mu_{Lavg}, s^{-1}\right)$ is equal to the maximum growth rate $(\mu_{max}, s^{-1})$ multiplied by the light intensity $\left(I_{ph}, mol_{ph}m^{-2}s^{-1}\right)$ and divided by the sum of the light intensity and of the light half-velocity constant $\left(K_L, mol_{ph}m^{-2}s^{-1}\right)$.

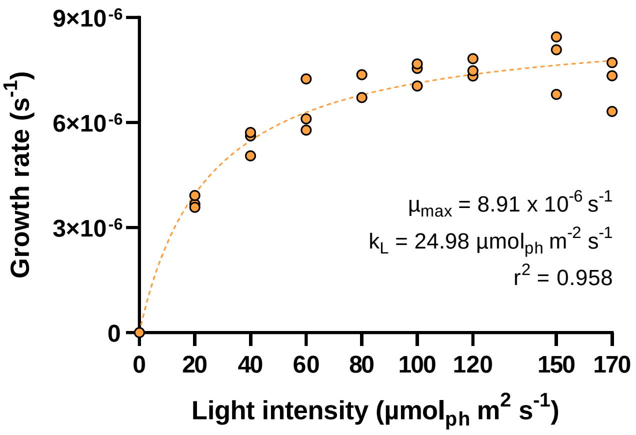


**Figure S3. Growth rate of *Anabaena* sp. PCC 7938 as a function of light intensity.** Dots represent experimental data and the dashed line, the fitted Monod equation.

**S7. Specific surface area of MGS-1 (BET)**

The specific surface area of MGS-1 (purchased from Exolith Lab, Orlando, Florida; shipped May 2019) was quantified by volumetric dinitrogen gas adsorption experiments with a BELSORP-mini (BEL Japan, Inc., Japan) apparatus followed by BET (Brunauer-Emmett-Teller) analysis. Prior to dinitrogen adsorption, samples were incubated at 120°C for at least 3 h under vacuum (< 2 mbar) and subsequently exposed to an argon atmosphere at room temperature for 30 min.

### S8. Equivalent system mass analysis

#### S8.1 Mass estimation

The overall mass of the system is the sum of the masses of the photobioreactor structure $(Reactor_{kg})$, sensors, actuators, and controllers $(Control_{kg})$, illumination system $(Illumination_{kg})$ and pumps $(Pumps_{kg})$:

$$M = Reactor_{kg} + Control_{kg} + Illumination_{kg} + Pumps_{kg} \qquad \text{(SE11)}$$

All mass values are in kilograms.

The mass of the photobioreactor PMMA's structure is pressure-dependent and is determined as described previously (*14*), except for the addition of an inner cylinder serving as a riser (1.53-m tall and of a diameter such that the riser-to-downcomer cross sectional area ratio is equal to 2). The illumination system is composed of two subsystems. One is providing natural light, collected at the Martian surface and delivered to the photobioreactor via fibre optics. This subsystem is assumed to have a mass of 18.7 kg $m^{-3}$, which corresponds to that estimated for one squared meter of crop production (*62*). The second light subsystem provides artificial light via LED lamps which are placed in the outer shell irradiating inwards, and in the inner shell irradiating in- and outwards. We assume an average of 4.5 high-pressure sodium lamps (0.21 kg each) per square meter (*56*), over an area corresponding to that of the outer shell plus twice that of the inner shell (see Section S7.2). The photobioreactors are expected to be equipped with a variety of actuators (e.g., valves), controllers and sensors related to, for instance, pressure, flow rates, temperature, pH, gas composition, and light intensity. We assume a combined mass for this category of 3 kg per cubic meter of culture. We also assume two liquid pumps weighing 1.5 kg each, and one air pump of 1 kg, per cubic meter of culture.

#### S8.2. Power requirements

The total power consumption of the photobioreactor is the sum of that from the sensors, actuators, and controllers $(Control_E)$, illumination system $(Illumination_E)$ and pumps $(Pumps_E)$:

$$P = Control_E + Illumination_E + Pumps_E \qquad \text{(SE12)}$$

All power values are expressed in kilowatt.

The power consumption of the illumination subsystem providing natural light is considered negligible. That of the subsystem providing artificial light is calculated as follows:

$$Illumination_{SystemE} = \frac{mean\ (I_{ph0})}{LED_E} \times \left(2\pi \times Base_r \times H_{culture} + \left(2 \times 2\pi \times \frac{Base_r}{\sqrt{2.7}} \times H_{culture} \times 0.9\right)\right) \times 0.001 \times Ratio_{LEDSunlight} \quad \text{(SE13)}$$

In Equation SE13, the average incident light intensity $\left(mean\ (I_{ph0}), mol_{ph}m^{-2}s^{-1}\right)$ is divided by the conversion factor between LED light intensity and Joule $\left(LED_E,\ mol_{ph}\,J^{-1}\right)$(*56*). It is then multiplied by the total area of incident light, which is composed of the outer shell cylinder area $(2\pi \times Base_r \times H_{culture},\ m^2)$ and twice the inner shell cylinder area $\left(2 \times 2\pi \times \frac{Base_r}{\sqrt{3}} \times H_{culture} \times 0.9, m^2\right)$. The inner shell is 90% as high as the outer shell. Its radial position is calculated assuming that the riser contains approximately one third of the photobioreactor volume. The watt value is then converted to kilowatt. Finally, a factor is applied which corresponds to the ratio between the provided artificial and natural light $\left(Ratio_{LEDSunlight}\right)$. This ratio is estimated at 0.5455, assuming that artificial light is active about 50% of the time on clear days and during dust storms (≈ 50 every 550 days)(*63*). Sensors, controllers and actuators have an estimated power consumption of 0.02 kW per cubic meter of culture.

The power requirement of liquid pumps is estimated at 25 W per pump. That of the air pump is assessed based on the oxygen removal power requirement of fine bubble diffusers (0.5 kWh $kg_{O2}^{-1}$) (*64*).

#### S8.3 Cost of consumables

Water-related costs were assessed based on estimates made for a system that relies on excavated regolith yielding 1.3% water at 300°C, in an early mission (*65*). That system has a mass of ≈ 1,400 kg $\left(Water_{systemkg}\right)$ and a power use of ≈ 45 kW $\left(Water_{systemE}\right)$. This includes the excavator rover that gathers regolith. The system yields 43,070 kg of water over a 480-day period $\left(Water_{systemp}\right)$. The assumed volume of this system is 33.6157 m$^3$ $\left(Water_{systemV}\right)$, calculated based on the volume-to-mass ratio provided in Table 6-11 of the addendum to the DRA 5.0 (*63*), assuming a 3-wt% water content and a 24-hour operation period. No cooling or crew time are considered for the water extraction system. The cost of water $(Water_{cost}, kg\ kg_{water}^{-1})$ is calculated as follows:

$$Water_{cost} = \frac{Water_{systemkg} + Water_{systemE} \times P_{eq} + Water_{systemV} \times V_{eq}}{Water_{systemp}} \quad \text{(SE14)}$$

Costs associated with water mining are assumed to also cover regolith extraction costs, as ca. 77 kilograms of regolith are processed to obtain one kilogram of water.

Carbon dioxide is assumed to be recovered from a continuously-operated atmospheric acquisition subsystem with a mass of 492.12 kg $\left(CarbonDioxide_{systemkg}\right)$, a power use of 17.863 kW $(CarbonDioxide_{systemE})$, and a volume of 0.66 m$^3$ $\left(CarbonDioxide_{systemV}\right)$. A carbon dioxide mass of 35,192 kg is produced $\left(CarbonDioxide_{systemp}\right)$(*63*). No cooling or crew time are considered for the atmospheric acquisition system. The cost of carbon dioxide $\left(CarbonDioxide_{cost}, kg\ kg_{CO_2}^{-1}\right)$ is calculated as follows:

$$CarbonDioxide_{cost} = \frac{CarbonDioxide_{systemkg} + CarbonDioxide_{systemE} \times P_{eq} + CarbonDioxide_{systemV} \times V_{eq}}{CarbonDioxide_{systemp}} \quad \text{(SE15)}$$

Nitrogen-enriched air is assumed to be a byproduct of carbon dioxide purification, and not to be further processed to remove argon or trace gases.
